# Protective activity of mRNA vaccines against ancestral and variant SARS-CoV-2 strains

**DOI:** 10.1101/2021.08.25.457693

**Authors:** Baoling Ying, Bradley Whitener, Laura A. VanBlargan, Ahmed O. Hassan, Swathi Shrihari, Chieh-Yu Liang, Courtney E. Karl, Samantha Mackin, Rita E. Chen, Natasha M. Kafai, Samuel H. Wilks, Derek J. Smith, Juan Manuel Carreño, Gagandeep Singh, Florian Krammer, Andrea Carfi, Sayda Elbashir, Darin K. Edwards, Larissa B. Thackray, Michael S. Diamond

## Abstract

Although mRNA vaccines prevent COVID-19, variants jeopardize their efficacy as immunity wanes. Here, we assessed the immunogenicity and protective activity of historical (mRNA-1273, designed for Wuhan-1 spike) or modified (mRNA-1273.351, designed for B.1.351 spike) preclinical Moderna mRNA vaccines in 129S2 and K18-hACE2 mice. Immunization with high or low dose formulations of mRNA vaccines induced neutralizing antibodies in serum against ancestral SARS-CoV-2 and several variants, although levels were lower particularly against the B.1.617.2 (Delta) virus. Protection against weight loss and lung pathology was observed with all high-dose vaccines against all viruses. Nonetheless, low-dose formulations of the vaccines, which produced lower magnitude antibody and T cell responses, and serve as a possible model for waning immunity, showed breakthrough lung infection and pneumonia with B.1.617.2. Thus, as levels of immunity induced by mRNA vaccines decline, breakthrough infection and disease likely will occur with some SARS-CoV-2 variants, suggesting a need for additional booster regimens.

## INTRODUCTION

Severe acute respiratory syndrome coronavirus 2 (SARS-CoV-2) is the cause of the Coronavirus Disease 2019 (COVID-19) syndrome. More than 211 million infections and 4.4 million deaths have been recorded worldwide (https://covid19.who.int) since the start of the pandemic. The extensive morbidity and mortality associated with the COVID-19 pandemic made the development of SARS-CoV-2 vaccines a global health priority. In a short period of less than one year, several highly effective vaccines targeting the SARS-CoV-2 spike protein encompassing multiple platforms (lipid nanoparticle encapsulated mRNA, inactivated virion, or viral-vectored vaccine platforms (Graham, 2020)) gained Emergency Use Authorization or Food and Drug Administration approval and were deployed with hundreds of millions of doses given worldwide (https://covid19.who.int). The currently used vaccines all were designed against the spike protein of strains that were circulating early in the pandemic. In localities with high rates of vaccination, markedly reduced numbers of infections, hospitalizations, and deaths were initially observed.

Despite the success of COVID-19 vaccines and their potential for curtailing the pandemic, the continued evolution of more transmissible SARS-CoV-2 variants of concern (VOC) including B.1.1.7 (Alpha), B.1.351 (Beta), B.1.1.28 (Gamma), and B.1.617.2 (Delta) with substitutions in the spike protein jeopardizes the efficacy of vaccination campaigns (Krause et al., 2021). Experiments in cell culture suggest that neutralization by vaccine-induced sera is diminished against variants expressing mutations in the spike gene at positions L452, E484, and elsewhere (Chen et al., 2021b; McCallum et al., 2021a; Tada et al., 2021; Wang et al., 2021a; Wang et al., 2021b; Wibmer et al., 2021). Moreover, viral-vectored (ChAdOx1 nCoV-19 and Ad26.CoV2) and protein nanoparticle (NVX-CoV2373)-based vaccines showed reduced activity (10 to 60%) against symptomatic infection caused by the B.1.351 variant in clinical trials in humans (Madhi et al., 2021; Sadoff et al., 2021; Shinde et al., 2021), whereas mRNA-based vaccines (*e.g*., BNT162b2) retained substantial (∼75%) efficacy against the B.1.351 variant in humans with almost complete protection against severe disease (Abu-Raddad et al., 2021).

Immunization of humans with two 100 μg doses of the lipid nanoparticle-encapsulated mRNA-1273 vaccine encoding a proline-stabilized full-length SARS-CoV-2 spike glycoprotein corresponding to the historical Wuhan-Hu-1 virus conferred 94% efficacy against symptomatic COVID-19 in clinical trials performed in the United States (Baden et al., 2021). More recent data in non-human primates shows that vaccination with two doses of mRNA-1273 results in an effective immune response that controls upper and lower respiratory tract infection after challenge with the SARS-CoV-2 B.1.351 viral variant (Corbett et al., 2021). As an alternative approach, several manufacturers have designed modified vaccines that target specific VOC including B.1.351 for possible immunization or boosting. Indeed, a mRNA-1273.351 vaccine recently was generated, which encodes a proline stabilized full-length SARS-CoV-2 spike glycoprotein from the B.1.351 virus. Here, we evaluated the immunogenicity and protective activity of lipid-encapsulated mRNA-1273 and mRNA-1273.351 Moderna vaccines in the context of challenge of wild-type 129S2 and human ACE2 (hACE2) transgenic (K18-hACE2) mice with historical and emerging SARS-CoV-2 strains including several key VOC.

## RESULTS

### Immunogenicity of mRNA vaccines in 129S2 mice

We first tested preclinical versions of the Moderna mRNA-1273 and mRNA-1273.351 vaccines encoding sequenced-optimized prefusion-stabilized spike proteins of Wuhan-1 and B.1.351, respectively, in immunocompetent 129S2 mice. These animals are permissive to infection by some SARS-CoV-2 variants (*e.g*., B.1.1.7, B.1.1.28, and B.1.351) or mouse-adapted strains (Chen et al., 2021a; Gu et al., 2020; Rathnasinghe et al., 2021) that encode an N501Y mutation, which enables engagement of endogenous murine ACE2 (Liu et al., 2021b). Infection of 129S2 mice with SARS-CoV-2 results in mild to moderate lung infection and clinical disease with subsequent recovery (Chen et al., 2021a; Rathnasinghe et al., 2021). To assess the immunogenicity of the vaccines, groups of 7 to 9-week-old female 129S2 mice were immunized and boosted three weeks later by an intramuscular route with 5 μg (high) or 0.25 μg (low) doses of mRNA-1273, mRNA-1273.351, mRNA-1273.211 (1:1 mixture [total 5 or 0.25 μg] of mRNA-1273 and mRNA-1273.351), or a control non-coding mRNA (**Fig 1A**); we included the mRNA-1273.211 mixture since it is being tested in humans (NCT04927065 (Wu et al., 2021)). Serum samples were collected three weeks after boosting, and IgG responses against recombinant spike proteins of ancestral (Wuhan-1) or variant (B.1.1.7, B.1.351, or B.1.617.2) viruses (Amanat et al., 2021) were evaluated by ELISA (**Fig 1B**). As expected, the control mRNA did not generate spike-specific IgG (values below the limit of detection), whereas antibody responses against the spike proteins from all other mRNA vaccines were robust. For the 5 μg dose, mean endpoint titers of serum ranged from 619,650 to 1,503,560 against the different spike proteins with little variation between the mRNA vaccines. For the 0.25 μg dose, approximately 5-fold lower serum IgG responses were observed with mean endpoint titers ranging from 126,900 to 382,725, again with little difference between the mRNA vaccines. The responses trended slightly higher against the ancestral spike protein and lower against the B.1.617.2 spike protein although most of these differences did not attain statistical significance. Overall, both doses and all spike-based vaccine formulations generated strong anti-spike protein IgG responses in 129S2 mice.

**Figure 1.**
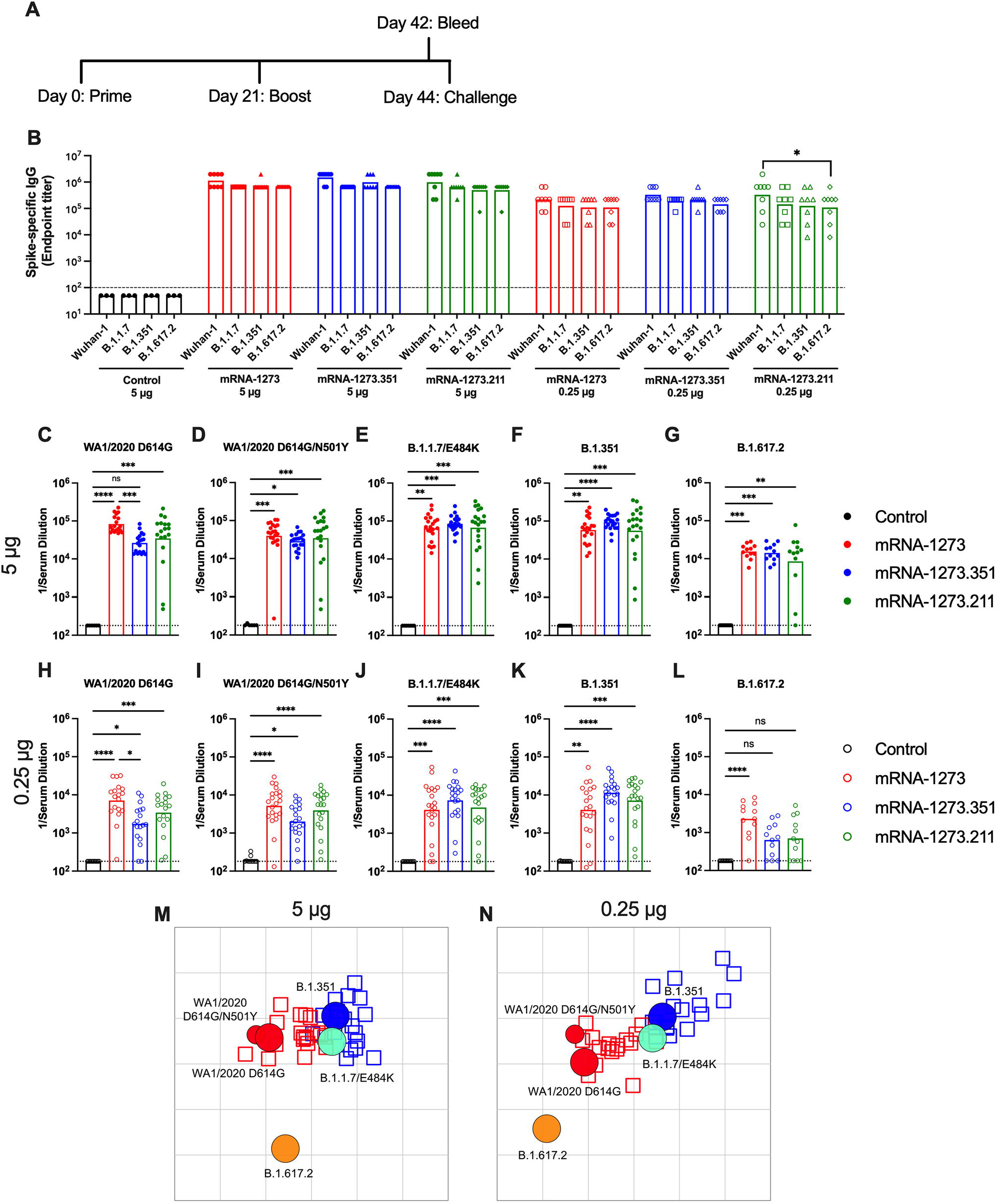
Immunogenicity analysis of mRNA vaccines in 129S2 mice. Seven to nine-week-old female 129S2 mice were immunized and boosted with 5 or 0.25 μg of mRNA vaccines. **A.** Scheme of immunizations, blood draw, and virus challenge. **B**. Serum anti-spike IgG responses at three weeks after booster immunization with mRNA vaccines (control (black symbols), mRNA-1273 (red symbols), mRNA-1273.351 (blue symbols), and mRNA-1273.211 (green symbols) against indicated spike proteins (Wuhan-1, B.1.1.7, B.1.351, or B.1.617.2) (n = 3 (control vaccine) or 8 (spike vaccines), two independent experiments, boxes illustrate mean values, dotted line shows the limit of detection (LOD); two-way ANOVA with Tukey’s post-test: *, *P* < 0.05). **C-L**. Serum neutralizing antibody responses three weeks after boosting as assessed by FRNT (half-maximal reduction, FRNT_50_ values) with WA1/2020 D614G (**C, H**), WA1/2020 D614G/N501Y (**D, I**), B.1.1.7/E484K (**E, J**), B.1.351 (**F, K**), or B.1.617.2 (**G, L**) in mice immunized with 5 (**C-G**) or 0.25 (**H-L**) μg of control (n = 6-10), mRNA-1273, mRNA-1273.351, or mRNA-1273.211 (n = 12-21) vaccines (two independent experiments, boxes illustrate geometric mean values, dotted line shows LOD; one-way Kruskal-Wallis ANOVA with Dunn’s post-test: *, *P* < 0.05; **, *P* < 0.01; ***, *P* < 0.001; **** *P* < 0.0001). **M-N.** Antigenic map of sera from 129S2 mice titrated against WA1/2020 D614G, WA1/2020 N501Y/D614G, B.1.1.7/E484K, B.1.351, and B.1.617.2. The maps show sera from mice that received 5 μg (**M**) or 0.25 μg (**N**) doses, respectively. Antigens (viruses) are shown as circles (WA1/2020 D614G: red, bigger circle, WA1/2020 N501Y/D614G: red, smaller circle, B.1.1.7/E484K: turquoise, B.1.351: blue, and B.1.617.2: orange) and sera as squares (blue for mRNA-1273.351-induced sera and red for mRNA-1273-induced sera). The X and Y axes correspond to antigenic distance, with one grid line corresponding to a two-fold serum dilution in the neutralization assay. The antigens and sera are arranged on the map such that the distances between them best represent the distances measured in the neutralization assay.

We characterized serum antibody responses functionally by assaying inhibition of SARS-CoV-2 infectivity using a focus-reduction neutralization test (FRNT) (Case et al., 2020b). We tested a panel of sera from each group of vaccinated mice against several fully-infectious SARS-CoV-2 strains including an ancestral Washington strain with a single D614G substitution (WA1/2020 D614G) or one with both D614G and N501Y substitutions (WA1/2020 D614G/N501Y), a B.1.1.7 isolate encoding an E484K mutation (B.1.1.7/E484K), a B.1.351 isolate, and a B.1.617.2 isolate (**Fig 1C-L**). Due to the limited amount of sera recovered from live animals, we started dilutions at 1/180. As expected, serum from all control mRNA-immunized mice did not inhibit infection of the SARS-CoV-2 strains (**Fig 1C-L**). For the 5 μg dose, all three spike gene vaccines (mRNA-1273, mRNA-1273.351, and mRNA-1273.211) induced robust serum neutralizing antibody responses (**Fig 1C-G**). In general, these titers were similar with the exception of ∼4-fold lower geometric mean titers (GMTs) against WA1/2020 D614G and ∼2-fold higher GMTs against B.1.351 induced by mRNA-1273.351 compared to the mRNA-1273 and mRNA-1273.211 vaccines. Lower neutralizing responses (∼4- to 5-fold) were seen against the B.1.617.2 strain by all three mRNA vaccines (**Fig 1G**). For the 0.25 μg vaccine dose, we observed ∼10-fold lower levels of serum neutralizing activity against each of the viruses (**Fig 1H-L**) and noted the following trends: (a) the mRNA-1273.351 vaccine induced lower levels of neutralizing antibody against WA1/2020 D614G and WA1/2020 D614G/N501Y than the mRNA-1273 vaccine (**Fig 1H-I**); (b) the mRNA-1273.211 mixture generally induced neutralizing antibodies that were equivalent to one of the two vaccine components; (c) serum from mRNA-1273-vaccinated mice showed smaller reductions in neutralization against B.1.351 than anticipated based on prior studies in humans and C57BL6 mice (Chen et al., 2021b; Wang et al., 2021a) (**Fig 1J**); (d) serum from mRNA-1273.351-vaccinated mice trended toward higher neutralization against B.1.351; and (e) serum neutralizing antibody levels from all vaccinated mice were lower against B.1.617.2 than other strains, although responses from animals administered mRNA-1273 were somewhat higher (**Fig 1L**). Overall, these differences were visualized best in a comparative analysis of the inhibitory activity of each serum sample for the 5 μg (**Fig S1A-C**) and 0.25 μg (**Fig S1D-F**) doses.

Using the neutralization data from mRNA vaccinated 129S2 mice, we created antigenic maps to visualize the relationships between the WA1/2020 D614G, WA1/2020 D614G/N501Y, B.1.1.7/E484K, B.1.351, and B.1.617.2 SARS-CoV-2 strains (**Fig 1M-N**). Neutralization titers obtained after 5 or 0.25 μg dosing with mRNA-1273 and mRNA-1273.351 vaccines were used to position the serum relative to each virus using antigenic cartography (a modification of multidimensional scaling for binding assay data), such that higher neutralization titers are represented by shorter distances between serum and the virus. Each gridline, or antigenic unit, of the map corresponds to a 2-fold difference in neutralization titer of a given virus. Three antigen clusters were observed: (a) WA1/2020 D614G and WA1/2020 D614G/N501Y grouped together; (b) viruses containing E484K mutations (B.1.1.7/E484K and B.1.351) had a similar antigenic position; and (c) B.1.617.2 was the most distant antigenically, which is consistent with the lower levels of serum neutralization induced by all of the mRNA vaccines against this VOC.

### Protection by mRNA vaccines in 129S2 mice

We tested the protective activity of the different mRNA vaccines in 129S2 mice. Three weeks after boosting, mice were challenged via an intranasal route with WA1/2020 N501Y/D614G, B.1.1.7/E484K, or B.1.351. The WA1/2020 D614G and B.1.617.2 viruses were not used for challenge in this model since they lack the mouse-adapting N501Y substitution and cannot infect conventional laboratory mice (Gu et al., 2020). Compared to the control mRNA vaccine, the 5 μg or 0.25 μg doses of mRNA-1273, mRNA-1273.351, or mRNA-1273.211 vaccines all prevented weight loss between 2 and 4 dpi, although protection was not statistically significant for some groups immunized with the mRNA-1273 vaccine and challenged with B.1.351 or B.1.1.7/E484K viruses (**Fig 2A-B**).

**Figure 2.**
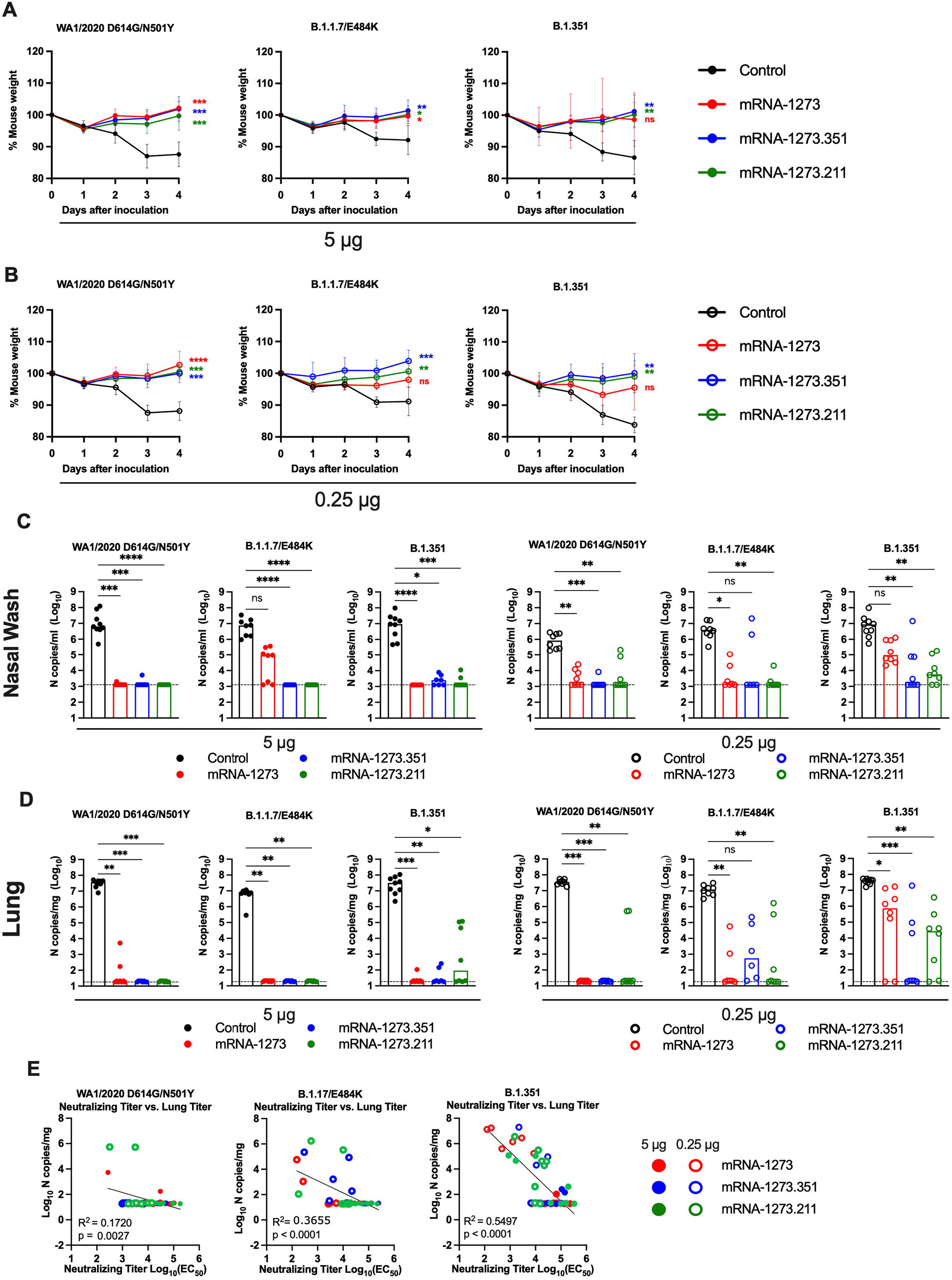
Protection against SARS-CoV-2 infection after mRNA vaccination in 129S2 mice. Seven to nine-week-old female 129S2 mice were immunized and boosted with 5 or 0.25 μg of mRNA vaccines (control (black symbols), mRNA-1273 (red symbols), mRNA-1273.351 (blue symbols), and mRNA-1273.211 [mixture, green symbols]) as described in Figure 1A. Three weeks after boosting, mice were challenged via intranasal inoculation with 10^5^ focus-forming units (FFU) of WA1/2020 N501Y/D614G, B.1.1.7/E484K, or B.1.351. **A-B**. Body weight change over time. Data shown is the mean +/− SEM (n = 6-9, two independent experiments; one-way ANOVA of area under the curve from 2-4 dpi with Dunnett’s post-test, comparison to control immunized group: ns, not significant; *P* > 0.05; *, *P* < 0.05; **, *P* < 0.01; ***, *P* < 0.001; **** *P* < 0.0001). **C-D**. Viral burden at 4 dpi in the nasal washes (**C**) and lungs (**D**) as assessed by qRT-PCR of the *N* gene after challenge of immunized mice with the indicated mRNA vaccines (n = 6-8, two independent experiments, boxes illustrate median values, dotted line shows LOD; one-way Kruskal-Wallis ANOVA with Dunn’s post-test, comparison among all immunization groups: ns, not significant; *P* > 0.05; *, *P* < 0.05; **, *P* < 0.01; ***, *P* < 0.001; **** *P* < 0.0001). **E**. Correlation analyses comparing serum neutralizing antibody concentrations three weeks after boosting plotted against lung viral titer (4 dpi) in 129S2 mice after challenge with the indicated SARS-CoV-2 strain (Pearson’s correlation *P* and R^2^ values are indicated as insets; closed symbols 5 μg vaccine dose; open symbols, 0.25 μg vaccine dose).

At 4 days post-infection (dpi), mice were euthanized, and nasal washes, lungs, and spleen were collected for viral burden analysis. In the nasal washes or lungs from control mRNA-vaccinated 129S2 mice, high levels (∼10^7^ copies of *N* per mL or mg) of viral RNA were measured after challenge with WA1/2020 N501Y/D614G, B.1.1.7/E484K, or B.1.351 (**Fig 2C-D**). Lower levels of SARS-CoV-2 RNA (∼10^2^ to 10^4^ copies of *N* per mg) were measured in the spleen (**Fig S2A**). In general, the mRNA-1273, mRNA-1273.351, and the mRNA-1273.211 vaccines conferred robust protection against infection in nasal washes, lungs, and spleens by the challenge SARS-CoV-2 strains, although some breakthrough was noted. After the 5 μg dose immunization with mRNA-1273, moderate B.1.1.7/E484K infection was detected in nasal washes in 5 of 8 mice, although viral RNA was absent from the lungs. Some (3 of 8) mice immunized with the mRNA-1273.211 mixture also showed breakthrough in the lungs, albeit at greater than 100-fold lower levels than the control vaccine. In comparison, the 5 μg dose of mRNA-1273.351 was protective in the nasal wash and lungs against all viruses, with little, if any, viral RNA measured.

As expected, the 0.25 μg dose of the mRNA vaccines showed less protective efficacy against SARS-CoV-2 challenge. With few exceptions (2 mice, mRNA-1273.351), the protection conferred by the 0.25 μg dose against WA1/2020 N501Y/D614G and B.1.1.7/E484K challenge was strong in the nasal washes at 4 dpi (**Fig 2C**). In comparison, after B.1.351 challenge, 8 of 8 mice immunized with mRNA-1273 showed viral RNA in nasal washes, with 3 of 8 showing levels that approached those seen in control-vaccinated mice. Greater protection was generated against B.1.351 by mRNA-1273.351 or the mRNA-1273.211 mixture vaccine, although breakthrough infections were detected. In the lungs, strong protection against infection with WA1/2020 N501Y/D614G was generated by all three spike mRNA vaccines (**Fig 2D**). However, some infection was seen after B.1.1.7/E484K or B.1.351 challenge. For example, 6 of 8 mice immunized with mRNA-1273 had moderate to high levels of B.1.351 viral RNA in their lungs at 4 dpi.

We assessed for correlations between vaccine-induced neutralizing antibody titers and protection against SARS-CoV-2 infection in the lung after virus challenge. Serum levels of neutralizing antibody were associated inversely with SARS-CoV-2 RNA levels in the lung (**Fig 2E**) with a minimum neutralizing titer of approximately 5,000 required to prevent infection in the lung at 4 dpi. Most of the breakthrough infections occurred with the B.1.351 challenge at the 0.25 μg dose of vaccines. For reasons that remains unclear, the threshold for complete protection in the lung after challenge with WA1/2020 N501Y/D614G was lower (2 to 7-fold) than against the other viruses. Moreover, when we compared body weight change at 4 dpi with neutralizing titers, only animals challenged with B.1.351 showed a linear correlation (**Fig S2B**), possibly because of the greater number of breakthrough infections in this group.

We also assessed the effect of the mRNA vaccines on lung disease at 4 dpi in129S2 mice. For these studies, we analyzed lung sections from the group of mice that received the lower 0.25 μg vaccine dose and the B.1.351 challenge virus, as this combination resulted in the greatest number of breakthrough infections. As expected, mice immunized with the control mRNA vaccine and challenged with B.1.351 developed mild pneumonia characterized by immune cell accumulation in perivascular and alveolar locations, vascular congestion, and interstitial edema. In contrast, animals immunized with mRNA-1273, mRNA-1273.351, or mRNA-1273.211 vaccines did not show these pathological changes (**Fig 3**). Thus, immunization with even the low dose of the mRNA vaccines was sufficient to mitigate SARS-CoV-2-induced lung injury in immunocompetent 129S2 mice challenged with some VOC.

**Figure 3.**
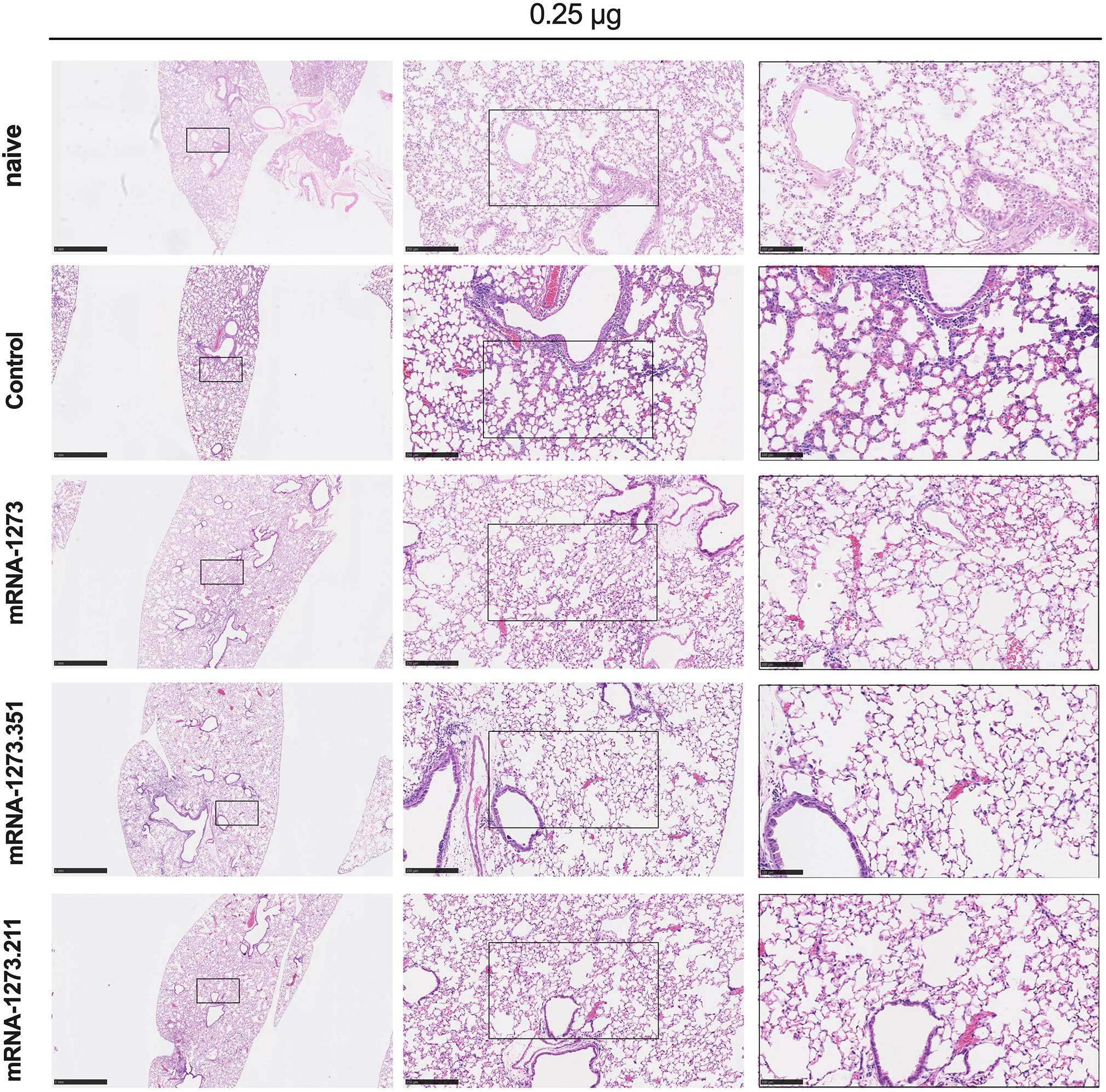
Vaccine-mediated protection against lung pathology in 129S2 mice. Seven to nine-week-old female 129S2 mice were immunized, boosted with 0.25 μg of mRNA vaccines (control, mRNA-1273, mRNA-1273.351, or mRNA-1273.211), and challenged with B.1.351 as described in Figure 2. Hematoxylin and eosin staining of lung sections harvested at 4 dpi or from a mock-infected animal. Images show low- (top; scale bars, 1 mm) and high-power (bottom; scale bars, 50 μm). Representative images from n = 2 per group.

### Immunogenicity of mRNA vaccines in K18-hACE2 transgenic mice

We next evaluated the mRNA-1273 and mRNA-1273.351 vaccines in K18-hACE2 transgenic mice, which sustain higher levels of infection and disease after intranasal inoculation by many SARS-CoV-2 strains (Winkler et al., 2020) including isolates containing or lacking mouse-adapting mutations (*e.g*., N501Y) (Chen et al., 2021a). Due to a limited availability of K18-hACE2 mice and the need to test two control viruses (WA1/2021 D614G and WA1/2021 D614G/N501Y), we tested mRNA-1273 and mRNA-1273.351 but not the mRNA-1273.211 mixture vaccine. Groups of 7-week-old female K18-hACE2 mice were immunized and boosted three weeks later by intramuscular route with 5 or 0.25 μg doses of mRNA-1273, mRNA-1273.351, or control mRNA vaccine (**Fig 4A**). Serum samples were collected three weeks after boosting, and IgG responses against recombinant spike proteins (Wuhan-1, B.1.1.7, B.1.351, or B.1.617.2) were evaluated by ELISA (**Fig 4B**). Antibody responses against the different spike proteins were robust although slightly lower (∼2 to 3-fold) than that seen in 129S2 mice (**Fig 1B**). Serum mean endpoint IgG titers ranged from 218,700 to 1,601,425 against the different spike proteins with little variation observed with the 5 μg doses of different mRNA vaccines. For the 0.25 μg dose, lower (∼6 to 10-fold) IgG titers were measured (24,300 to 101,250) with little difference between the mRNA-1273 and mRNA-1273.351 vaccines. Although the IgG levels against the B.1.617.2 spike protein were reduced slightly compared to the other SARS-CoV-2 spike proteins, in general, robust antibody responses were detected in K18-hACE2 mice.

**Figure 4.**
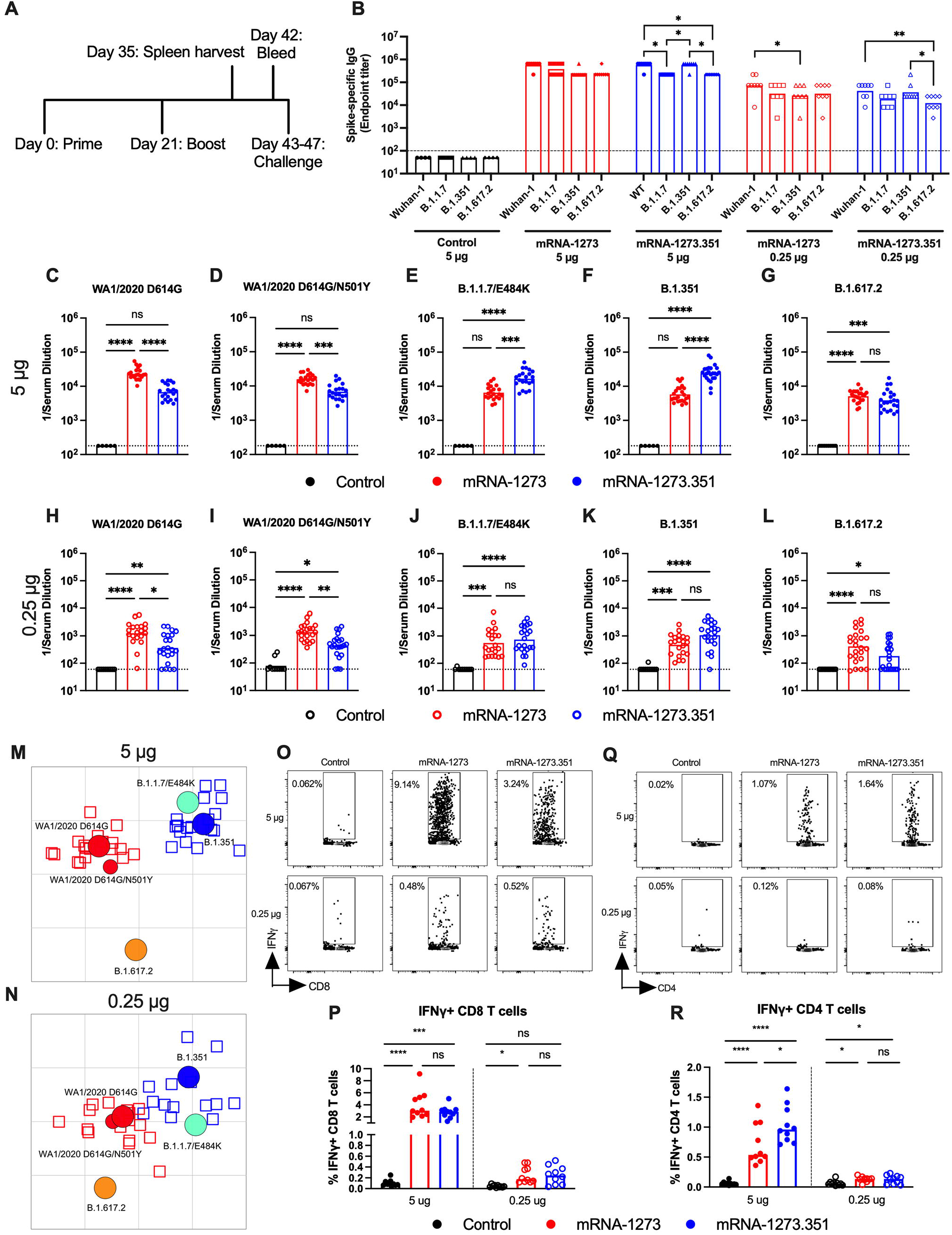
Immunogenicity analysis of mRNA vaccines in K18-hACE2 transgenic mice. Seven-week-old female K18-hACE2 mice were immunized and boosted with 5 or 0.25 μg of mRNA vaccines (control (black symbols), mRNA-1273 (red symbols), or mRNA-1273.351 (blue symbols). **A.** Scheme of immunizations, spleen harvest, blood draw, and virus challenge. **B**. Serum anti-spike IgG responses at three weeks after booster immunization with mRNA vaccines (control (black symbols), mRNA-1273 (red symbols), mRNA-1273.351 (blue symbols) against indicated spike proteins (Wuhan-1, B.1.1.7, B.1.351, or B.1.617.2). (n = 3 (control vaccine) or 8 (spike vaccines), two independent experiments, boxes illustrate mean value, dotted line shows the LOD; two-way ANOVA with Tukey’s post-test: *, *P* < 0.05; **, *P* < 0.01). **C-L**. Serum neutralizing antibody responses three weeks after boosting as judged by FRNT (half-maximal reduction, FRNT_50_ values) with WA1/2020 D614G (**C, H**), WA1/2020 D614G/N501Y (**D, I**), B.1.1.7/E484K (**E, J**), B.1.351 (**F, K**), or B.1.617.2 (**G, L**) in mice immunized with 5 (**C-G**) or 0.25 (**H-L**) μg of control (n = 5-10), mRNA-1273 (n = 20-24), and mRNA-1273.351 (n = 21-24) vaccines (two independent experiments, boxes illustrate geometric mean values, dotted line shows LOD; one-way Kruskal-Wallis ANOVA with Dunn’s post-test: *, *P* < 0.05; **, *P* < 0.01; ***, *P* < 0.001; **** *P* < 0.0001). **M-N.** Antigenic map of sera from K18-hACE2 mice titrated against WA1/2020 D614G, WA1/2020 N501Y/D614G, B.1.1.7/E484K, B.1.351, and B.1.617.2. The maps show sera from immunized mice that received 5 μg (**M**) or 0.25 μg (**N**) doses, respectively with symbol details described in Figure 1. **O-R**. CD8^+^ (**O, P**) and CD4^+^ (**Q, R**) T cell responses in K18-hACE2 mice at day 35, two weeks after booster immunization with mRNA vaccines (n= 10 for each group, two independent experiments, boxes illustrate median values, one-way ANOVA with Tukey’s post-test: ns, not significant; *, *P* < 0.05; **, *P* > 0.01; ***, *P* < 0.001; **** *P* < 0.0001).

We performed FRNTs to assess the neutralizing activity of pre-challenge serum against WA1/2020 D614G, WA1/2020 D614G/N501Y, B.1.1.7/E484K, B.1.351, and B.1.617.2 SARS-CoV-2 strains. Because of the limited amount of sera recovered from K18-hACE2 mice, we initially started dilutions at 1/180. As expected, serum from all control mRNA-immunized mice did not inhibit infection of the SARS-CoV-2 strains (**Fig 4C-L**). In general, neutralizing antibody titers induced by 5 or 0.25 μg mRNA vaccine dosing trended lower (∼ 3 to 6-fold) in immunized K18-hACE2 than from 129S2 mice. For the 5 μg dose, while both mRNA-1273 and mRNA-1273.351 vaccines induced robust serum neutralizing antibody responses, we observed the following (**Fig 4C-G and Fig S3**): (a) the mRNA-1273.351 vaccine induced lower levels of neutralizing antibody against WA1/2020 D614G and WA1/2020 D614G/N501Y than the mRNA-1273 vaccine (**Fig 4C and D**); (b) a reciprocal pattern was observed against viruses containing E484K mutations. The mRNA-1273.351 vaccine induced higher levels of neutralizing antibody against B.1.1.7/E484K and B.1.351 than the mRNA-1273 vaccine (**Fig 4E and F**); and (c) no differences in neutralizing activity were observed with the mRNA-1273 and mRNA-1273.351 vaccines against the B.1.617.2 strain. Although responses were elevated, they were lower than against other strains (**Fig 4G**). Similar patterns were observed for the 0.25 μg dose (**Fig 4H-L**), although ∼10-fold lower levels of neutralizing activity were induced by each vaccine against each of the viruses. Because of this, we started our dilution series at 1/60 for sera derived from animals immunized with the 0.25 μg dose of mRNA vaccines. In general, the pattern of neutralization paralleled results with the higher dose, with the mRNA-1273 vaccine performing better against historical WA1/2020 viruses and the mRNA-1273.351 vaccine showing greater inhibitory titers against B.1.351 (**Fig 4H, I and K**). However, serum from mice vaccinated with mRNA-1273 or mRNA-1273.351 vaccines neutralized B.1.617.2 less efficiently (**Fig 4L**), with several data points at the limit of detection (1/60: mRNA-1273, 4 of 24; mRNA-1273.351, 9 of 24) and responses induced by mRNA-1273 trending higher. A comparative analysis of the inhibitory activity of each serum sample for the 5 μg (**Fig S3A-B**) and 0.25 μg (**Fig SC-D**) doses visually showed these differences, as serum induced by the mRNA-1273 vaccine consistently showed less neutralizing activity against B.1.1.7/E484K, B.1.351, and B.1.617.2, whereas serum from mRNA-1273.351-vaccinated mice had greater inhibitory activity against B.1.351 and B.1.1.7/E484K.

We used the neutralization data from mRNA-vaccinated K18-hACE2 mice to generate maps defining the antigenic relationships between WA1/2020 D614G, WA1/2020 D614G/N501Y, B.1.1.7/E484K, B.1.351, and B.1.617.2 SARS-CoV-2 strains (**Fig 4M-N**). Serum obtained after 5 or 0.25 μg dosing with mRNA-1273 or mRNA-1273.351 vaccines was analyzed against the indicated viruses, and each antigenic unit corresponded to a 2-fold difference in neutralization titer of a given virus. The results were remarkably similar to that seen with 129S2 vaccinated mice (**Fig 1M-N**): (a) WA1/2020 D614G and WA1/2020 D614G/N501Y grouped together; (b) B.1.1.7/E484K and B.1.351 viruses, which contain E484K mutations, grouped near each other; and (c) B.1.617.2 localized to a separate antigenic group.

We also examined T cell responses in mRNA-vaccinated K18-hACE2 mice two weeks after boosting (**Fig 4A and M-Q**) using H-2^b^ restricted immunodominant peptides in the spike protein for CD8^+^ and CD4^+^ T cells. After peptide stimulation *ex vivo* and staining for intracellular IFN-γ production, we detected a robust CD8^+^ T cell (∼2 to 4 percent positive) response in the spleens of animals immunized with 5 μg of the mRNA-1273 or mRNA-1273.351 vaccines (**Fig 4O and P**). The response was approximately 10-fold lower in animals immunized with the lower 0.25 μg dose. While we also detected a spike protein-specific CD4^+^ T cell response after immunization (∼0.5 to 1.5 percent positive) with the 5 μg dose of mRNA-1273 or mRNA-1273.351 vaccines, it was lower in magnitude (**Fig 4Q and R**). Moreover, the low 0.25 μg dose mRNA-1273 or mRNA-1273.351 vaccines induced CD4^+^ T cell responses that were barely greater than the control mRNA vaccine.

### Protection by mRNA vaccines in K18-hACE2 transgenic mice

We next evaluated the protective activity of the mRNA vaccines in K18-hACE2 mice. Three to four weeks after boosting, mice were challenged via intranasal route with WA1/2020 D614G, WA1/2020 N501Y/D614G, B.1.1.7/E484K, B.1.351, or B.1.617.2 strains. Compared to the control mRNA vaccine, the 5 μg and 0.25 μg doses of mRNA-1273 and mRNA-1273.351 vaccines all prevented the weight loss occurring between 3 and 6 dpi (**Fig 5A-B**).

**Figure 5.**
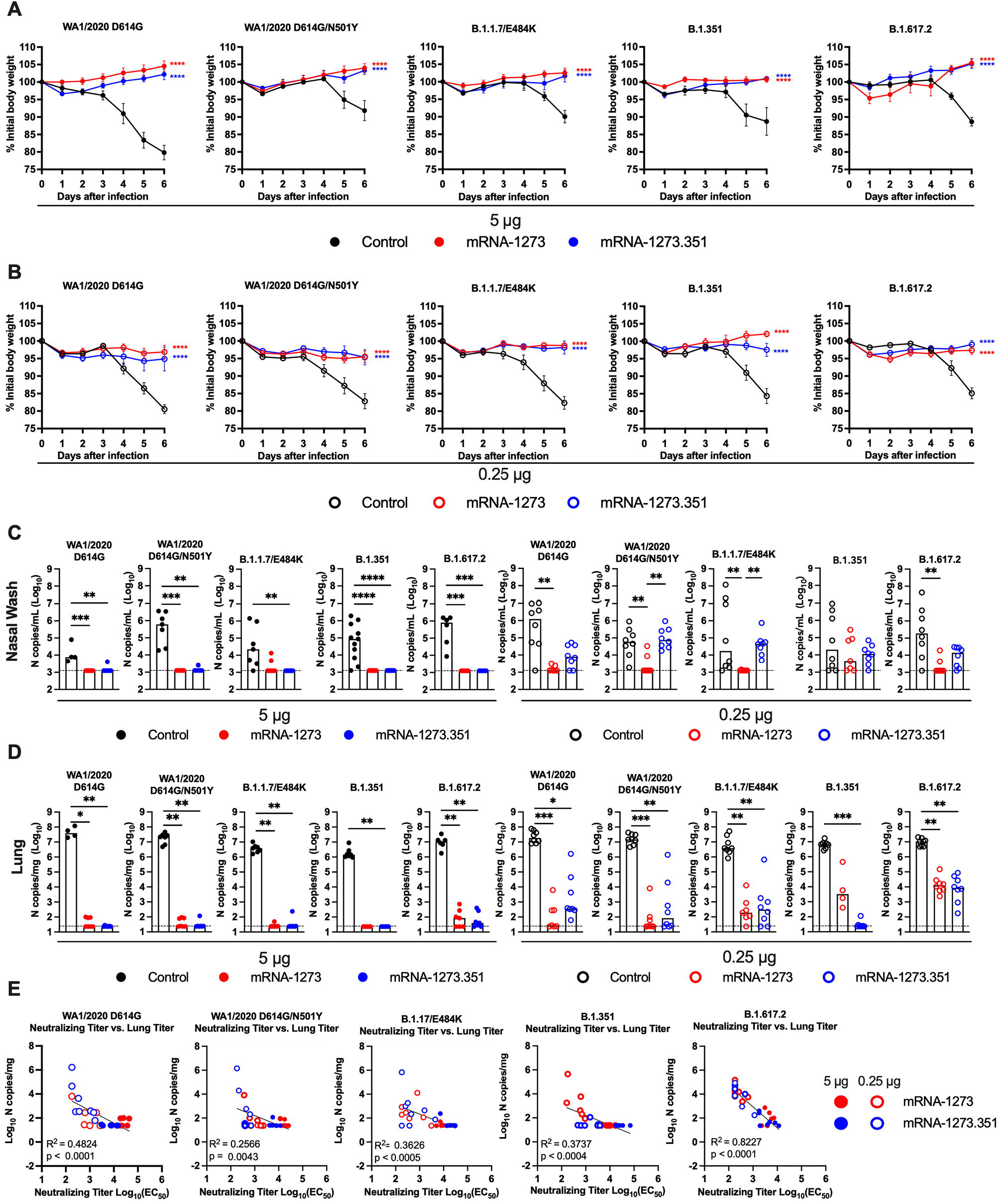
Protection against SARS-CoV-2 infection after mRNA vaccination in K18-hACE2 transgenic mice. Seven-week-old female K18-hACE2 mice were immunized and boosted with 5 or 0.25 μg of mRNA vaccines (control (black symbols), mRNA-1273 (red symbols), or mRNA-1273.351 (blue symbols) as described in Figure 4A. Four weeks after boosting, mice were challenged via intranasal inoculation with 10^3^ to 3 x 10^4^ FFU of WA1/2020 D614G, WA1/2020 N501Y/D614G, B.1.1.7/E484K, B.1.351, or B.1.617.2, depending on the strain. **A-B**. Body weight change over time. Data shown is the mean +/− SEM (n = 4-8, two independent experiments; one-way ANOVA of area under the curve from 2-4 dpi with Dunnett’s post-test, comparison to control immunized group: **** *P* < 0.0001). **C-D**. Viral burden levels at 6 dpi in the nasal washes (**C**) and lungs (**D**) as assessed by qRT-PCR of the *N* gene after challenge of immunized mice with the indicated mRNA vaccines (n = 4-8, two independent experiments, boxes illustrate median values, dotted line shows LOD; one-way Kruskal-Wallis ANOVA with Dunn’s post-test, comparison among all immunization groups: *, *P* < 0.05; **, *P* < 0.01; ***, *P* < 0.001; **** *P* < 0.0001). **E**. Correlation analyses comparing serum neutralizing antibody concentrations three weeks after boosting plotted against lung viral titer (6 dpi) in K18- hACE2 mice after challenge with the indicated SARS-CoV-2 strain (Pearson’s correlation *P* and R^2^ values are indicated as insets; closed symbols 5 μg vaccine dose; open symbols, 0.25 μg vaccine dose).

At 6 dpi, mice were euthanized, and nasal washes, lungs, and brains were collected for viral burden analysis (**Fig 5C-D, and S4A**). In the nasal washes of control mRNA-vaccinated K18-hACE2 mice, although some variability was observed, moderate levels (∼10^5^ copies of *N* per mL) of viral RNA were measured after challenge with WA1/2020 D614G, WA1/2020 N501Y/D614G, B.1.1.7/E484K, B.1.351, or B.1.617.2 strains (**Fig 5C**). In comparison, in the lungs of control mRNA-vaccinated K18-hACE2 mice, higher and more uniform levels (∼10^7^ copies of *N* per mg) of viral RNA were detected after challenge with all SARS-CoV-2 strains (**Fig 5D**). The brains of control RNA-vaccinated K18-hACE2 mice showed some variability, as seen previously (Winkler et al., 2020), with many but not all animals showing substantial infection (10^8^ copies of *N* per mL) (**Fig S4A**). The high 5 μg dose of mRNA-1273 or mRNA-1273.351 vaccines protected against infection in nasal washes, lung, and brain, with virtually no viral breakthrough regardless of the challenge strain. After the 0.25 μg dose immunization with mRNA-1273, a loss of protection against infection in the nasal washes (3 of 7 mice) and lungs (4 of 4 mice) was observed after challenge with B.1.351 and in the lungs only after challenge with B.1.1.7/E484K (6 of 7 mice) or B.1.617.2 (8 of 8 mice) viruses. After the 0.25 μg dose immunization with mRNA-1273.351, incomplete protection against infection in the nasal washes, lungs, and brain also was observed after challenge with WA1/2020 D614G (6, 7, and 4 of 8 mice, respectively), WA1/2020 D614G/N501Y (8, 4, and 6 of 8 mice, respectively), B.1.1.7/E484K (8, 6, and 3 of 8 mice, respectively), and B.1.617.2 (7, 8, and 5 of 8 mice, respectively). The 0.25 μg dose of the mRNA-1273.351 vaccine protected better against lung and brain infection by the homologous B.1.351 virus than against other strains.

We explored whether vaccine-induced neutralizing antibody titers correlated with protection after challenge with WA1/2020 D614G, WA1/2020 N501Y/D614G, B.1.1.7/E484K, B.1.351, or B.1.617.2 viruses. In general, serum levels of neutralizing antibody inversely correlated with viral RNA levels in the lung (**Fig 5E**) for all viruses, with more infection occurring in animals with lower neutralization titers. However, for WA1/2020 D614G, WA1/2020 N501Y/D614G, B.1.1.7/E484K, and B.1.351, some of the animals with low neutralization titers still were protected against infection in the lung. The correlation was most linear for B.1.617.2-challenged animals, with a minimum neutralizing titer of approximately 2,000 required to completely prevent infection at 6 dpi. Most of the breakthrough B.1.617.2 infections occurred with the 0.25 μg dose of mRNA vaccines. The threshold for complete protection in the lung after virus challenge varied somewhat with lower levels required for WA1/2020 D614G and WA1/2020 N501Y/D614G. When we compared body weight change in K18-hACE2 mice at 6 dpi with neutralizing titers, a linear relationship was observed with all challenge viruses except B.1.351 (**Fig S4B**). The best correlation was seen after B.1.617.2 challenge, with greater weight loss in mice immunized with the 0.25 μg vaccine dose and having lower serum neutralizing antibody titers.

Because a pro-inflammatory host response to SARS-CoV-2 infection can contribute to pulmonary pathology and severe COVID-19, we assessed the ability of the mRNA vaccines to suppress cytokine and chemokine levels in the lung after virus challenge (**Fig S5**). For these studies, K18-hACE2 mice were immunized and boosted with 5 or 0.25 μg of control, mRNA-1273 or mRNA-1273.351 vaccines and then challenged with WA1/2020 N501Y/D614G, B.1.351, or B.1.617.2. SARS-CoV-2 infection of control mRNA vaccinated K18-hACE2 mice resulted in high levels of expression in lung homogenates of several pro-inflammatory cytokines and chemokines including G-CSF, IFNγ, IL-1β, IL-6, CXCL1, CXCL5, CXCL9, CXCL10, CCL2, and CCL4. Pro-inflammatory cytokine and chemokines in the lung at 6 dpi generally were decreased in all mice vaccinated with 5 μg doses of mRNA-1273 or mRNA-1273.351 regardless of the challenge virus (**Fig S5A and B**). While this pattern also trended for the 0.25 μg dose of both mRNA vaccines, some cytokines and chemokines (e.g., IL-1β, IL-6, CXCL9, and CXCL10) remained elevated especially after challenge with B.1.617.2 (**Fig S5C and D**).

We evaluated the ability of the mRNA-1273 and mRNA-1273.351 vaccines to prevent disease in K18-hACE2 mice by performing histological analysis of lung tissues from immunized animals that were challenged with WA1/2020 D614G, WA1/2020 N501Y/D614G, B.1.1.7/E484K, B.1.351, or B.1.617.2. As expected, lung sections obtained at 6 dpi from mice immunized with the control mRNA vaccine and challenged with any of the SARS-CoV-2 strains showed severe pneumonia characterized by immune cell infiltration, alveolar space consolidation, vascular congestion, and interstitial edema (**Fig 6 and 7**). In comparison, mice immunized with the high 5 μg dose of mRNA-1273 or mRNA-1273.351 did not develop lung pathology, with histological findings similar to uninfected K18-hACE2 mice (**Fig 6**). Mice immunized with the low 0.25 μg dose of the mRNA vaccines however, showed different results (**Fig 7**): (a) mice vaccinated with mRNA-1273 showed few, if any, pathological changes after WA1/2020 D614G, WA1/2020 N501Y/D614G, or B.1.1.7/E484K challenge. Nonetheless, some mRNA-1273 vaccinated mice challenged with B.1.351 showed pulmonary vascular congestion and mild lung inflammation; (b) mice vaccinated with mRNA-1273.351 showed almost complete protection after WA1/2020 D614G, B.1.1.7/E484K, or B.1.351 challenge, whereas scattered inflammation and alveolar septal thickening was apparent in sections from some WA1/2020 N501Y/D614G challenged mice; (c) of note, lung sections from mice vaccinated with the lower 0.25 μg dose mRNA-1273 or mRNA-1273.351 and challenged with B.1.617.2 showed evidence of viral pneumonia with prominent foci of immune cells inflammation and airspace consolidation. Thus, low doses immunization of original or modified mRNA vaccines do not fully protect K18-hACE2 mice from challenge with B.1.617.2 and result in mild to moderate infection and lung pathology.

**Figure 6.**
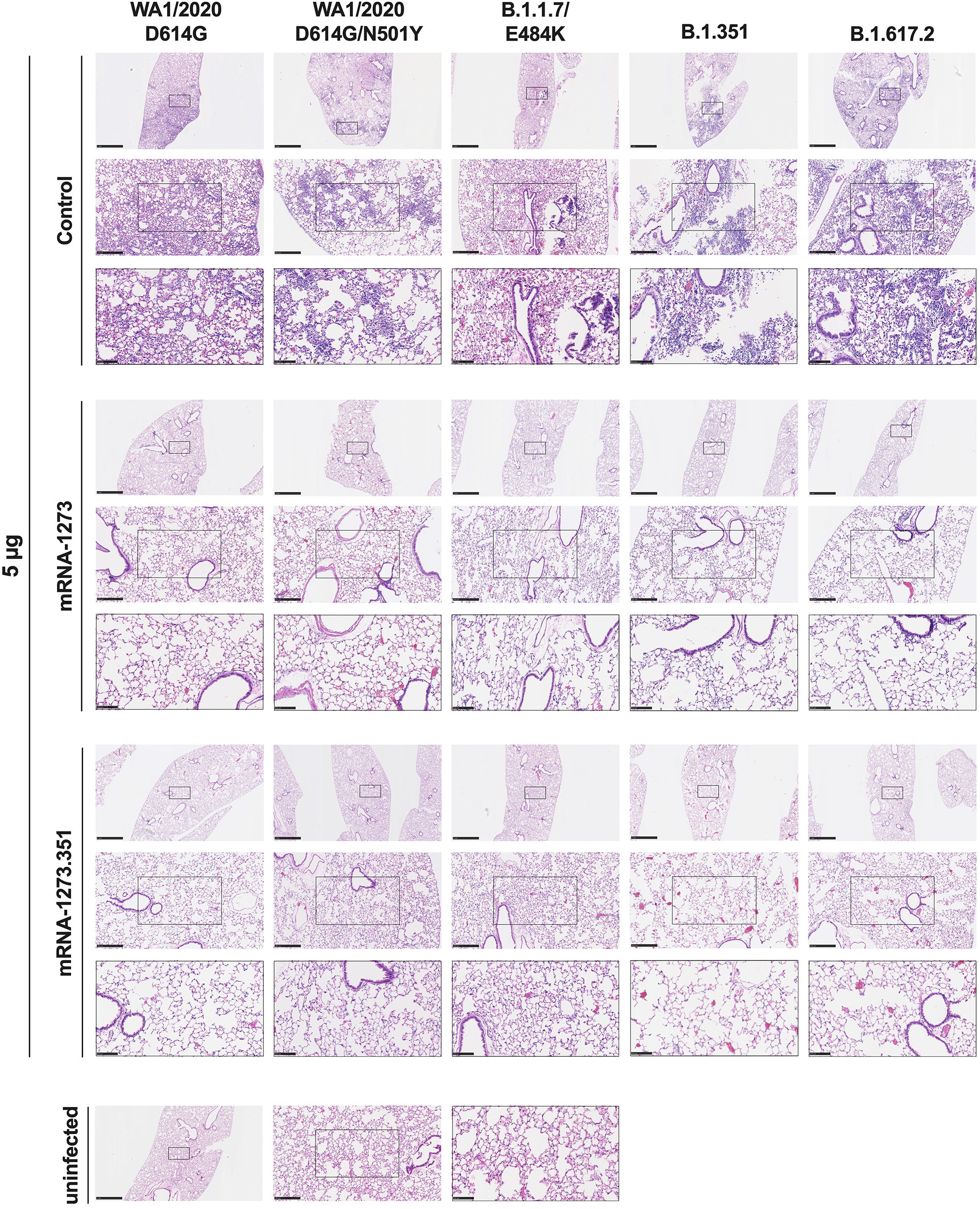
High dose mRNA vaccine protection against lung inflammation and pathology in K18-hACE2 transgenic mice. Seven to nine-week-old female K18-hACE2 transgenic mice were immunized, boosted with 5 μg of mRNA vaccines (control, mRNA-1273, or mRNA-1273.351, and challenged with WA1/2020, WA1/2020 N501Y/D614G, B.1.1.7/E484K, B.1.351, or B.1.617.2. as described in Figure 5. Hematoxylin and eosin staining of lung sections harvested at 6 dpi or from an uninfected animal. Images show low- (top; scale bars, 1 mm) and high-power (bottom; scale bars, 50 μm). Representative images of multiple lung sections from n = 2 per group.

**Figure 7.**
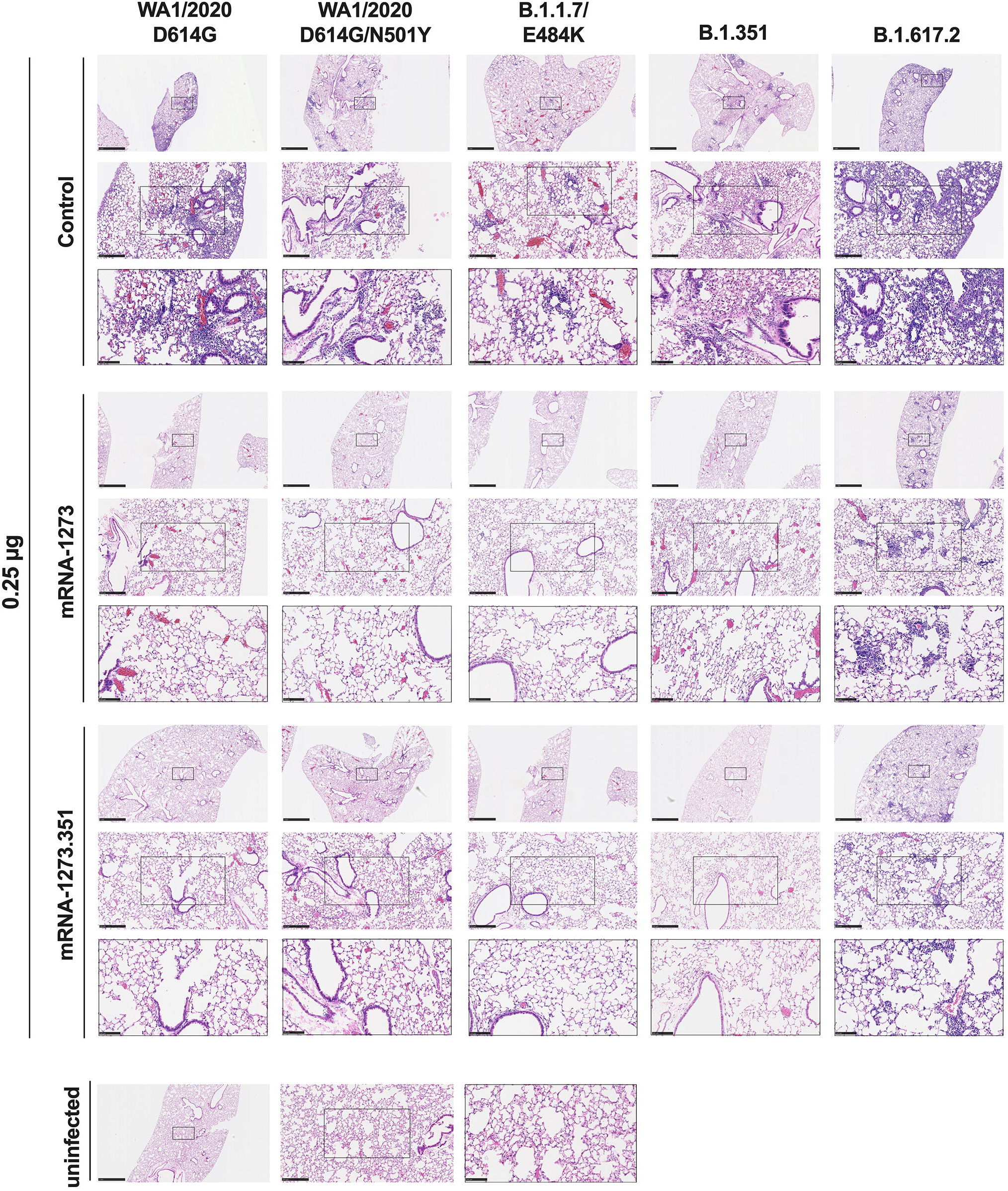
Low dose mRNA vaccine protection against lung inflammation and pathology in K18-hACE2 transgenic mice. Seven to nine-week-old female K18-hACE2 transgenic mice were immunized, boosted with 0.25 μg of mRNA vaccines (control, mRNA-1273, or mRNA-1273.351, and challenged with WA1/2020, WA1/2020 N501Y/D614G, B.1.1.7/E484K, B.1.351, or B.1.617.2. as described in Figure 5. Hematoxylin and eosin staining of lung sections harvested at 6 dpi or from an uninfected animal. Images show low- (top; scale bars, 1 mm) and high-power (bottom; scale bars, 50 μm). Representative images of multiple lung sections from n = 2 per group.

## DISCUSSION

Robust vaccine-induced immune responses and sustained protective activity against emerging SARS-CoV-2 variants are needed to limit human disease and curtail the COVID-19 global pandemic. A concern in the field is whether immunity generated by vaccines will wane sufficiently to lose activity against VOC with mutations or deletions in regions of the spike protein recognized by neutralizing antibodies. In the current study, we evaluated the immunogenicity and protective activity of high- and low-dose formulations of Moderna mRNA vaccines targeting historical (mRNA-1273) or variant (mRNA-1273.351) strains. The low-dose formulation study arm was designed to model waning immunity and assess for possible strain-specific breakthrough infections. Indeed, the lower neutralizing antibody and T cell responses measured with the 0.25 μg dose of mRNA-1273 or mRNA-1273.351 parallel those seen for human antibody and T cell responses six months after a primary vaccination series with mRNA-1273 (Mateus et al., 2021; Pegu et al., 2021).

Immunization of 129S2 or K18-hACE2 transgenic mice with mRNA-1273, mRNA-1273.351, or the mRNA-1273.211 mixture induced neutralizing antibodies against spike in serum against historical WA1/2020 and several key VOC. Challenge studies performed approximately one month after the second vaccine dose showed robust protection against weight loss and lung pathology with all high-dose vaccine formulations and infecting SARS-CoV-2 strains. Nonetheless, the low-dose vaccine formulations showed evidence of viral infection breakthrough and lung pathological changes consistent with pneumonia especially with the B.1.617.2 strain, which correlated with lower strain-specific neutralizing antibody levels. In general, variant-specific vaccine designs appeared to induce greater antibody responses and confer more protection against homologous virus strains.

Our experiments expand upon a preliminary immunogenicity study, which showed that vaccination of H-2^d^ BALB/c mice with mRNA-1273.351 resulted in high serum neutralizing antibody titers against the B.1.351 lineage, whereas the mRNA-1273.211 vaccine induced broad cross-variant neutralization (Wu et al., 2021). We performed experiments with two H-2^b^ strains, 129S2 and K18-hACE2 C57BL/6, and observed some similarities and differences. In K18-hACE2 mice, the mRNA-1273 vaccine, which encodes for the Wuhan-1 pre-fusion stabilized spike, induced higher neutralizing titers against WA1/2020 strains but lower responses against viruses containing E484K mutations in spike (B.1.1.7/E484K and B.1.351), which agrees with recent immunization studies in NHPs (Corbett et al., 2021). Reciprocally, the mRNA-1273.351 vaccine, which encodes for the B.1.351 pre-fusion stabilized spike, induced higher neutralizing titers against B.1.1.7/E484K and B.1.351. In 129S2 mice, only the mRNA-1273.351 vaccine induced a lower neutralizing response against WA1/2020 D614G, as the remainder of the neutralizing antibody responses were largely equivalent between vaccines. However, in both K18-hACE2 and 129S2 mice, the mRNA-1273 and mRNA-1273.351 vaccines induced antibody responses that neutralized B.1.617.2 less efficiently than the other SARS-CoV-2 strains. Analysis of serum antibodies and B cell repertoire against SARS-CoV-2 VOC from ongoing human clinical trials comparing mRNA-1273 and mRNA-1273.351 vaccines will be needed to corroborate our results obtained in small animal models. Indeed, the differences in neutralizing antibody titers induced by mRNA-1273 against WA1/2020 D614G and B.1.351 in mice were smaller in magnitude than that seen in humans one month after boosting, but similar to that observed six months after boosting (Pegu et al., 2021).

While the high-dose vaccination regimen with mRNA-1273 and mRNA-1273.351 induced robust neutralizing antibody and T cell responses that conferred almost complete protection against all strains, animals immunized with the low-dose scheme showed virological and pathological breakthrough that varied with the vaccine formulation and challenge strain. The low-dose vaccination approach we employed as a model for waning immunity resulted in approximately 10 to 20-fold reduced peak neutralization titers and T cell responses compared to the high-dose arm, which corresponds to the 90% loss observed 90 days after natural infection or vaccination in humans (Ibarrondo et al., 2021). The greatest loss in antibody neutralization (both 129S2 and K18-hACE2 mice) and protection (K18-hACE2 mice) consistently occurred with the B.1.617.2 variant. In contrast, recent longitudinal studies in humans immunized with mRNA-1273 showed lower levels of serum antibody recognition of B.1.351 than other VOC, although live virus neutralization assays were not performed with B.1.617.2 (Pegu et al., 2021). In our experiments with live virus, the loss of neutralizing activity was equivalent if not greater for B.1.617.2 than B.1.351, as seen by others (Liu et al., 2021a). Based on sequence changes in the spike protein (B.1.617.2: T19R, 156del, 157del, R158G, L452R, T478K, D614G, P681R, and D950N; and B.1.351: D80A, D215G, 241del, 242del, 243del, K417N, E484K, N501Y, D614G, A701V) and known binding sites in the receptor binding motif of neutralizing antibodies (at residue E484), it is not apparent why neutralizing activity and protection in mice were less against B.1.617.2 than B.1.351, although there was a direct correlation with levels of neutralizing antibody and B.1.617.2 burden in the lung. Nonetheless, mutations in the B.1.617.2 alter key antigenic sites and can abrogate recognition by neutralizing antibodies (McCallum et al., 2021b). Other possible explanations for the loss of potency of antibodies against B.1.617.2 include differential display of B.1.617.2 spike proteins on the surface of infected cells and engagement of Fc effector functions (Ravetch et al., 2021; Winkler et al., 2021) or differential ability of antibodies to block cell-to-cell spread in a strain-dependent manner (Kruglova et al., 2021). Our observation of B.1.617.2 infection and lung disease in low-dose mRNA-vaccinated K18-hACE2 mice as a model of waning immunity corresponds to descriptions of B.1.617.2 breakthrough infections in vaccinated humans in the United States, Israel, and elsewhere, some of which have required hospitalization (Brown et al., 2021; Puranik et al., 2021).

Our studies in 129S2 and K18-hACE2 mice with parental and modified mRNA vaccines show robust immunogenicity and protection against multiple SARS-CoV-2 strains when high-dose immunization schemes are used, although some differences in immunity are seen with particular vaccines against selected variants. While the lower dose of mRNA vaccines generally protected against matched virus challenge infection (e.g., mRNA-1273 vaccination and WA1/2020 challenge or mRNA-1273.351 vaccination and B.1.351 challenge), breakthrough events were seen with some non-matched challenges (e.g., mRNA-1273 vaccination and B.1.351 challenge or mRNA-1273.351 vaccination and WA1/2020 challenge). As the low dose of mRNA-1273 and 1273.351 vaccines induced lower neutralizing titers and protected less against challenge with the B.1.617.2 variant, higher titers will be needed to minimize B.1.617.2 infection, transmission, and disease. Although studies in humans are required, boosting with historical or variant (*e.g*., mRNA encoding B.1.617.2 spike genes) vaccines might be required to prevent breakthrough events as vaccine-induced immunity wanes.

### Limitations of study

We note several limitations in our study. (1) The studies in 129S2 mice precluded challenge with B.1.617.2, as it does not infect murine cells because it lacks an N501Y mutation. The generation of recombinant SARS-CoV-2 strains with spike genes encoding B.1.617.2 and an N501Y mutation could overcome this limitation. (2) Female 129S2 and K18-hACE2 mice were used to allow for group caging of the large cohorts required for these multi-arm vaccination studies. Follow-up experiments in male mice are needed to confirm results are not sex-biased. (3) We used lower vaccine dosing as a model for waning immunity. Studies that directly address durability of immune responses and protection are needed for corroboration. (4) We used historical, variant, or mixed mRNA vaccine formulations with homologous boosting schemes. Animals studies that test heterologous boosting (mRNA-1273 prime followed by mRNA-1273.351 boost) (Wu et al., 2021) also are needed to support clinical trials. (5) Our studies focused on immunogenicity and protection in two strains of mice because of the ability to set up large animal cohorts and the tools available for analysis. These results require confirmation in other animal models of SARS-CoV-2 infection including hamsters and non-human primates (Muñoz-Fontela et al., 2020). (6) We did not establish direct immunological correlates of vaccine protection or failure for all vaccine and challenge strain pairs. While some relationships were more predictive (*e.g*., low B.1.617.2 neutralizing titers and viral burden in the lung), others were not.

## ACKNOWLEDGEMENTS

This study was supported by grants and contracts from NIH (R01 AI157155, HHSN75N93019C00074, T32 AI007172, NIAID Centers of Excellence for Influenza Research and Response (CEIRR) contracts HHSN272201400008C and 75N93021C00014, and the Collaborative Influenza Vaccine Innovation Centers (CIVIC) contract 75N93019C00051). This work also was supported by the Alafi Neuroimaging Laboratory, the Hope Center for Neurological Disorders, and NIH Shared Instrumentation Grant (S10 RR0227552). We thank Marciela DeGrace for help in study design and funding support, Richard Webby, Mehul Suthar, and Pei-Yong Shi for several of the viruses used in this study, Kristen Valentine and Sujan Shresta for providing immunodominant T cell peptides in advance of publication, Barbara Mühlemann for help with antigenic cartography, and Emma Winkler and Oleksandr Dmytrenko for providing lung sections from naive mice. We acknowledge the Pulmonary Morphology Core at Washington University School of Medicine for tissue sectioning and slide imaging.

## AUTHOR CONTRIBUTIONS

L.A.V. and B.Y. performed and analyzed neutralization assays. B.Y., B.W., A.O.H., L.A.V., S.S., C.E.K., S.M., N.M.K., R.E.C., and L.B.T. performed mouse experiments. B.W., S.S., and C-Y.L. performed viral burden analyses. A.O.H. and B.Y. performed T cell analyses. J.M.C., G.S., and F.K. performed and analyzed the ELISA experiments. S.H.W and D.J.S. performed the antigenic cartography analysis. A.C., S.E., and D.E. provided the mRNA vaccines and helped design experiments. M.S.D. obtained funding. L.B.T. and M.S.D. supervised the research. M.S.D. and L.B.T. wrote the initial draft, with the other authors providing editorial comments.

## SUPPLEMENTAL FIGURE LEGENDS

**Figure S1. Analysis of serum neutralization of SARS-CoV-2 strains from 129S2 mice immunized with mRNA vaccines, Related to Figure 1.** Comparison of neutralizing activity of sera against WA1/2020 D614G, WA1/2020 D614G/N501Y, B.1.1.7/E484K, B.1.351, and B.1.617.2. Sera was obtained three weeks after boosting with 5 μg (**A-C**) or 0.25 μg (**D-F**) of mRNA vaccines: mRNA-1273 (**A and D**), mRNA-1273.351 (**B and E**), and mRNA-1273.211 (**C and F**). Results are from experiments performed in **Figure 1C-L**. Geometric mean neutralization titers (GMT) are shown above each graph, dotted line represents the LOD. Solid lines connect data points from the same serum sample across strains.

**Figure S2. Protection against SARS-CoV-2 infection after mRNA vaccination in 129S2 mice, Related to Figure 2**. Seven to nine-week-old female 129S2 mice were immunized and boosted with 5 or 0.25 μg of mRNA vaccines as described in **Figure 1A.** Three weeks after boosting, mice were challenged via intranasal inoculation with 10^5^ FFU of WA1/2020 N501Y/D614G, B.1.1.7/E484K, or B.1.351. **A.** Viral burden at 4 dpi in the spleen as assessed by qRT-PCR of the *N* gene after challenge of immunized mice with the indicated mRNA vaccines (n = 6-8, two independent experiments; boxes illustrate median values, dotted line shows LOD; one-way Kruskal-Wallis ANOVA with Dunn’s post-test, comparison among all immunization groups: ns, not significant; *P* > 0.05; *, *P* < 0.05; **, *P* < 0.01; ***, *P* < 0.001; **** *P* < 0.0001). **B.** Correlation analyses comparing serum neutralizing antibody concentrations three weeks after boosting plotted against weight change in 129S2 mice after challenge with the indicated SARS-CoV-2 strain (Pearson’s correlation *P* and R^2^ values are indicated as insets; closed symbols, 5 μg vaccine dose; open symbols, 0.25 μg vaccine dose).

**Figure S3. Analysis of serum neutralization of SARS-CoV-2 strains from K18-hACE2 mice immunized with mRNA vaccines, Related to Figure 4.** Comparison of neutralizing activity of sera against WA1/2020 D614G, WA1/2020 D614G/N501Y, B.1.1.7/E484K, B.1.351, and B.1.617.2. Sera was obtained three weeks after boosting with 5 μg (**A-B)** or 0.25 μg (**C-D**) mRNA vaccines: mRNA-1273 (**A and C**) and mRNA-1273.351 (**B and D**), Results are from experiments performed in **Figure 4C-L**. GMTs are shown above each graph, dotted line represents the LOD. Solid lines connect data points from the same serum sample across strains.

**Figure S4. Protection against SARS-CoV-2 infection after mRNA vaccination in K18-hACE2 transgenic mice, Related to Figure 5**. Seven-week-old K18-hACE2 mice were immunized and boosted with 5 or 0.25 μg of mRNA vaccines (control (black symbols), mRNA-1273 (red symbols), or mRNA-1273.351 (blue symbols) as described in **Figure 4A.** Three to four weeks after the last boost, mice were challenged via intranasal inoculation with WA1/2020 D614, WA1/2020 N501Y/D614G, B.1.1.7/E484K, B.1.351, or B.1.617.2 as described in **Figure 5**. **(A)** Viral burden levels at 6 dpi in the brain as assessed by qRT-PCR of the *N* gene after challenge of immunized mice with the indicated mRNA vaccines (n = 4-8, two independent experiments; boxes illustrate median values, dotted line shows LOD; one-way Kruskal-Wallis ANOVA with Dunn’s post-test, comparison among all immunization groups: *, *P* < 0.05; **, *P* < 0.01; ***, *P* < 0.001; **** *P* < 0.0001). **B**. Correlation analyses comparing serum neutralizing antibody concentrations three weeks after boosting plotted against weight change in K18-hACE2 mice after challenge with the indicated SARS-CoV-2 strain (Pearson’s correlation *P* and R^2^ values are indicated as insets; closed symbols, 5 μg vaccine dose; open symbols, 0.25 μg vaccine dose).

**Figure S5. Cytokine induction in lungs after mRNA vaccination and SARS-CoV-2 challenge, Related to Figure 5.** Cytokine levels from mice immunized with 5 μg (**A-B**) or 0.25 μg (**C-D**) dose of mRNA vaccines as measured by multiplex platform in lung tissues of SARS-CoV-2-infected mice at 6 dpi. **A and C**. For each cytokine, fold-change was calculated compared to mock-inoculated mice and log_2_ (fold-change) was plotted in the corresponding color-coded heat-map. **B and D**. Cytokine levels as measured by multiplex platform in the lungs of SARS-CoV-2-infected mice after vaccination (n = 7-8 per group, two independent experiments, mean values +/− SEM is shown).

## STAR METHODS

### RESOURCE AVAILABLITY

#### Lead Contact

Further information and requests for resources and reagents should be directed to and will be fulfilled by the Lead Contact, Michael S. Diamond.

#### Materials Availability

All requests for resources and reagents should be directed to and will be fulfilled by the Lead Contact author. This includes mice, antibodies, viruses, vaccines, proteins, and peptides. All reagents will be made available on request after completion of a Materials Transfer Agreement.

#### Data and code availability

All data supporting the findings of this study are available within the paper and are available from the corresponding author upon request.

### EXPERIMENTAL MODEL AND SUBJECT DETAILS

#### Cells

Vero-TMPRSS2 (Zang et al., 2020) and Vero-hACE2-TMPRRS2 (Chen et al., 2021b) cells were cultured at 37°C in Dulbecco’s Modified Eagle medium (DMEM) supplemented with 10% fetal bovine serum (FBS), 10□mM HEPES pH 7.3, 1□mM sodium pyruvate, 1× non-essential amino acids, and 100□U/mL of penicillin–streptomycin. Vero-TMPRSS2 cells were supplemented with 5 μg/mL of blasticidin. Vero-hACE2-TMPRSS2 cells were supplemented with 10 µg/mL of puromycin. All cells routinely tested negative for mycoplasma using a PCR-based assay.

#### Viruses

The WA1/2020 recombinant strain with substitutions (D614G or N501Y/D614G) were obtained from an infectious cDNA clone of the 2019n-CoV/USA_WA1/2020 strain as described previously (Plante et al., 2020). The B.1.351, B.1.1.7/E484K, and B.1.617.2 isolates were originally obtained from nasopharyngeal isolates. All viruses were passaged once in Vero-TMPRSS2 cells and subjected to next-generation sequencing as described previously (Chen et al., 2021b) to confirm the introduction and stability of substitutions. All virus experiments were performed in an approved biosafety level 3 (BSL-3) facility.

#### Mice

Animal studies were carried out in accordance with the recommendations in the Guide for the Care and Use of Laboratory Animals of the National Institutes of Health. The protocols were approved by the Institutional Animal Care and Use Committee at the Washington University School of Medicine (assurance number A3381–01). Virus inoculations were performed under anesthesia that was induced and maintained with ketamine hydrochloride and xylazine, and all efforts were made to minimize animal suffering.

Heterozygous K18-hACE2 C57BL/6J mice (strain: 2B6.Cg-Tg(K18-ACE2)2Prlmn/J) and 129 mice (strain: 129S2/SvPasCrl) were obtained from The Jackson Laboratory and Charles River Laboratories, respectively. Animals were housed in groups and fed standard chow diets.

#### Pre-clinical vaccine mRNA and lipid nanoparticle production process

A sequence-optimized mRNA encoding prefusion-stabilized Wuhan-Hu-1 (mRNA-1273) or B.1.351-variant (mRNA-1273.351) SARS-CoV-2 S-2P protein was synthesized *in vitro* using an optimized T7 RNA polymerase-mediated transcription reaction with complete replacement of uridine by N1m-pseudouridine (Nelson et al., 2020). The reaction included a DNA template containing the immunogen open-reading frame flanked by 5’ untranslated region (UTR) and 3’ UTR sequences and was terminated by an encoded polyA tail. After transcription, the cap-1 structure was added to the 5’ end using the vaccinia virus capping enzyme (New England Biolabs) and vaccinia virus 2’-O-methyltransferase (New England Biolabs). The mRNA was purified by oligo-dT affinity purification, buffer exchanged by tangential flow filtration into sodium acetate, pH 5.0, sterile filtered, and kept frozen at –20°C until further use.

The mRNA was encapsulated in a lipid nanoparticle through a modified ethanol-drop nanoprecipitation process described previously (Hassett et al., 2019). Ionizable, structural, helper, and polyethylene glycol lipids were briefly mixed with mRNA in an acetate buffer, pH 5.0, at a ratio of 2.5:1 (lipid:mRNA). The mixture was neutralized with Tris-HCl, pH 7.5, sucrose was added as a cryoprotectant, and the final solution was sterile-filtered. Vials were filled with formulated lipid nanonparticle and stored frozen at –20°C until further use. The vaccine product underwent analytical characterization, which included the determination of particle size and polydispersity, encapsulation, mRNA purity, double-stranded RNA content, osmolality, pH, endotoxin, and bioburden, and the material was deemed acceptable for *in vivo* study.

#### Antigens

Recombinant soluble S proteins from different SARS-CoV-2 strains were expressed as previously described (Amanat et al., 2021; Stadlbauer et al., 2020). Briefly, mammalian cell codon-optimized nucleotide sequences coding for the soluble ectodomain of the S protein of SARS-CoV-2 including a C-terminal thrombin cleavage site, T4 foldon trimerization domain, and hexahistidine tag were cloned into mammalian expression vector pCAGGS. The spike protein sequence was modified to remove the polybasic cleavage site (RRAR to A), and two pre-fusion stabilizing proline mutations were introduced (K986P and V987P, wild type Wuhan-Hu-1 numbering). Recombinant proteins were produced in Expi293F cells (ThermoFisher) by transfection with purified DNA using the ExpiFectamine 293 Transfection Kit (ThermoFisher). Supernatants from transfected cells were harvested 3 days post-transfection, and recombinant proteins were purified using Ni-NTA agarose (ThermoFisher), then buffer exchanged into PBS and concentrated using Amicon Ultracel centrifugal filters (EMD Millipore).

### METHOD DETAILS

#### ELISA

Assays were performed in 96-well microtiter plates (Thermo Fisher) coated with 50 µL of recombinant spike from wild type SARS-CoV-2 or variant viruses B.1.1.7, B.1.351, or B.1.617.2. Plates were incubated at 4°C overnight and then blocked with 200 µL of 3% non-fat dry milk (AmericanBio) in PBS containing 0.1% Tween-20 (PBST) for one hour at room temperature (RT). Sera were serially diluted in 1% non-fat dry milk in PBST and added to the plates. Plates were incubated for 120 min at room temperature and then washed 3 times with PBST. Goat anti-mouse IgG-HRP (Sigma-Aldrich, 1:9000) was diluted in 1% non-fat dry milk in PBST before adding to the wells and incubating for 60 min at room temperature. Plates were washed 3 times with PBST before the addition of peroxidase substrate (SigmaFAST o-phenylenediamine dihydrochloride, Sigma-Aldrich). Reactions were stopped by the addition of 3 M hydrochloric acid. Optical density (OD) measurements were taken at 490 nm, and endpoint titers were calculated in excel using a 0.15 OD 490 nm cutoff. Graphs were generated using Graphpad Prism v9.

#### Focus reduction neutralization test

Serial dilutions of sera were incubated with 10^2^ focus-forming units (FFU) of different strains of SARS-CoV-2 for 1 h at 37°C. Antibody-virus complexes were added to Vero-TMPRSS2 cell monolayers in 96-well plates and incubated at 37°C for 1 h. Subsequently, cells were overlaid with 1% (w/v) methylcellulose in MEM. Plates were harvested 30 h later by removing overlays and fixed with 4% PFA in PBS for 20 min at room temperature. Plates were washed and sequentially incubated with an oligoclonal pool of SARS2-2, SARS2-11, SARS2-16, SARS2-31, SARS2-38, SARS2-57, and SARS2-71 (Liu et al., 2021c) anti-S antibodies and HRP-conjugated goat anti-mouse IgG (Sigma, 12-349) in PBS supplemented with 0.1% saponin and 0.1% bovine serum albumin. SARS-CoV-2-infected cell foci were visualized using TrueBlue peroxidase substrate (KPL) and quantitated on an ImmunoSpot microanalyzer (Cellular Technologies).

#### Mouse experiments

Female 129S2 (catalog 287) and K18-hACE2 C57BL/6 (catalog 034860) mice were purchased from the Charles River and The Jackson Laboratory, respectively. Seven to nine-week-old animals were immunized and boosted three weeks apart with 0.25 or 5 μg of mRNA vaccines (control, mRNA-1273, mRNA-1273.351, or mRNA-1273.211) in 50 µl PBS via intramuscular injection in the hind leg. Animals were bled at specified time points via the mandibular vein to obtain sera for immunogenicity analysis. Three to four weeks after boosting, mice were inoculated with 10^5^ FFU (129S2) or 10^3^ to 3 x 10^4^ FFU (K18-hACE2) of WA1/2020 D614G (10^4^), WA1/2020 N501Y/D614G (10^3^), B.1.1.7/E484K (10^3^), B.1.351 (10^3^), or B.1.617.2 (3 x 10^4^) of SARS-CoV-2 strains by the intranasal route. Different doses of viruses were used in K18-hACE2 mice to match weight loss and infection in the nasal wash and lungs. This approach was necessary as some viruses (WA1/2020 N501Y/D614G, B.1.1.7/E484K, and B.1.351) encode N501Y mutations that enhance pathogenicity in mice (Gu et al., 2020; Muruato et al., 2021; Rathnasinghe et al., 2021). Animals were euthanized at 4 or 6 dpi, and tissues were harvested for virological, immunological, and pathological analyses.

#### Measurement of viral burden

Tissues were weighed and homogenized with zirconia beads in a MagNA Lyser instrument (Roche Life Science) in 1000 μL of DMEM medium supplemented with 2% heat-inactivated FBS. Tissue homogenates were clarified by centrifugation at 10,000 rpm for 5 min and stored at −80°C. RNA was extracted using the MagMax mirVana Total RNA isolation kit (Thermo Fisher Scientific) on the Kingfisher Flex extraction robot (Thermo Fisher Scientific). RNA was reverse transcribed and amplified using the TaqMan RNA-to-CT 1-Step Kit (Thermo Fisher Scientific). Reverse transcription was carried out at 48°C for 15 min followed by 2 min at 95°C. Amplification was accomplished over 50 cycles as follows: 95°C for 15 s and 60°C for 1 min. Copies of SARS-CoV-2 *N* gene RNA in samples were determined using a previously published assay (Case et al., 2020a). Briefly, a TaqMan assay was designed to target a highly conserved region of the N gene (Forward primer: ATGCTGCAATCGTGCTACAA; Reverse primer: GACTGCCGCCTCTGCTC; Probe: /56-FAM/TCAAGGAAC/ZEN/AACATTGCCAA/3IABkFQ/). This region was included in an RNA standard to allow for copy number determination down to 10 copies per reaction. The reaction mixture contained final concentrations of primers and probe of 500 and 100 nM, respectively.

#### Cytokine and chemokine protein measurements

Lung homogenates were incubated with Triton-X-100 (1% final concentration) for 1 h at room temperature to inactivate SARS-CoV-2. Homogenates then were analyzed for cytokines and chemokines by Eve Technologies Corporation (Calgary, AB, Canada) using their Mouse Cytokine Array/Chemokine Array 31-Plex (MD31) platform.

#### Lung histology

Animals were euthanized before harvest and fixation of tissues. Lungs were inflated with ∼2 mL of 10% neutral buffered formalin using a 3-mL syringe and catheter inserted into the trachea and kept in fixative for 7 days. Tissues were embedded in paraffin, and sections were stained with hematoxylin and eosin. Images were captured using the Nanozoomer (Hamamatsu) at the Alafi Neuroimaging Core at Washington University.

#### Peptide restimulation and intracellular cytokine staining

Two weeks after boosting, splenocytes from vaccinated K18-hACE2 mice were stimulated *ex vivo* with an H-2D^b^-restricted CD8 or CD4 immunodominant peptide (amino acids 262–270 and 62-76 of the S protein, respectively; gift of K. Valentine and S. Shresta, La Jolla Institute for Immunology) for 16 h at 37°C with brefeldin A (BioLegend, 420601) added for the last 4 h of incubation. Following blocking with FcγR antibody (BioLegend, clone 93), cells were stained on ice with CD45 BUV395 (BD BioSciences clone 30-F11), CD4 PE (BD BioSciences clone GK1.5), CD8 FITC (BioLegend clone 53-6.7), and Fixable Aqua Dead Cell Stain (Invitrogen, L34966). Stained cells were fixed and permeabilized with the Foxp3/Transcription Factor Staining Buffer Set (eBiosciences, 00-5523). Subsequently, intracellular staining was performed with anti-IFN-γ Alexa 647 (BioLegend, clone XMG1.2), and anti-TNFα BV605 (BioLegend, clone MP6-XT22). Analysis was performed on a BD LSRFortessa X-20 cytometer, using FlowJo X 10.0 software.

#### Antigenic cartography

A target distance from an individual serum to each virus was derived by calculating the difference between the logarithm (log2) reciprocal neutralization titer for that particular virus and the log2 reciprocal maximum titer achieved by that serum (against any virus). Thus, the higher the reciprocal titer, the shorter the target distance. As the log2 of the reciprocal titer was used, a 2-fold change in titer equates to a fixed change in target distance whatever the magnitude of the actual titers. Antigenic cartography (Smith et al., 2004) then was used to optimize the positions of the viruses and sera relative to each other on a map, minimizing the sum-squared error between map distance and target distance. Each virus is therefore positioned by multiple sera, and the sera themselves also are positioned only by their distances to the viruses. Hence, sera with different neutralization profiles to the virus panel are in separate locations on the map but contribute equally to positioning of the viruses.

### QUANTIFICATION AND STATISTICAL ANALYSIS

Statistical significance was assigned when *P* values were < 0.05 using Prism Version 10 (GraphPad). Tests, number of animals (n), median values, and statistical comparison groups are indicated in the Figure legends.

