## Supplementary figures and images for "Protective activity of mRNA vaccines against ancestral and variant SARS-CoV-2 strains"

### Figure S1

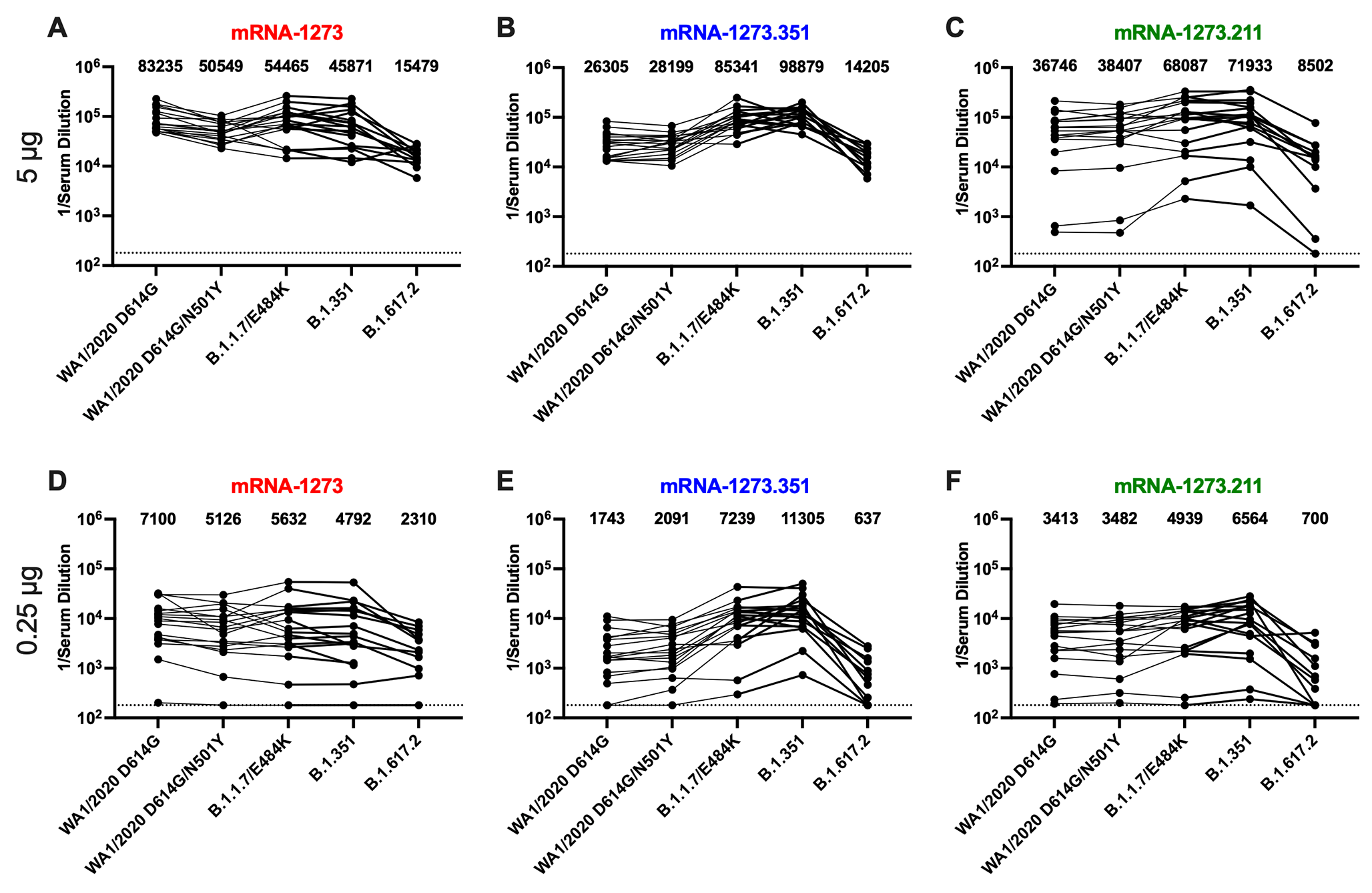

### Figure S2

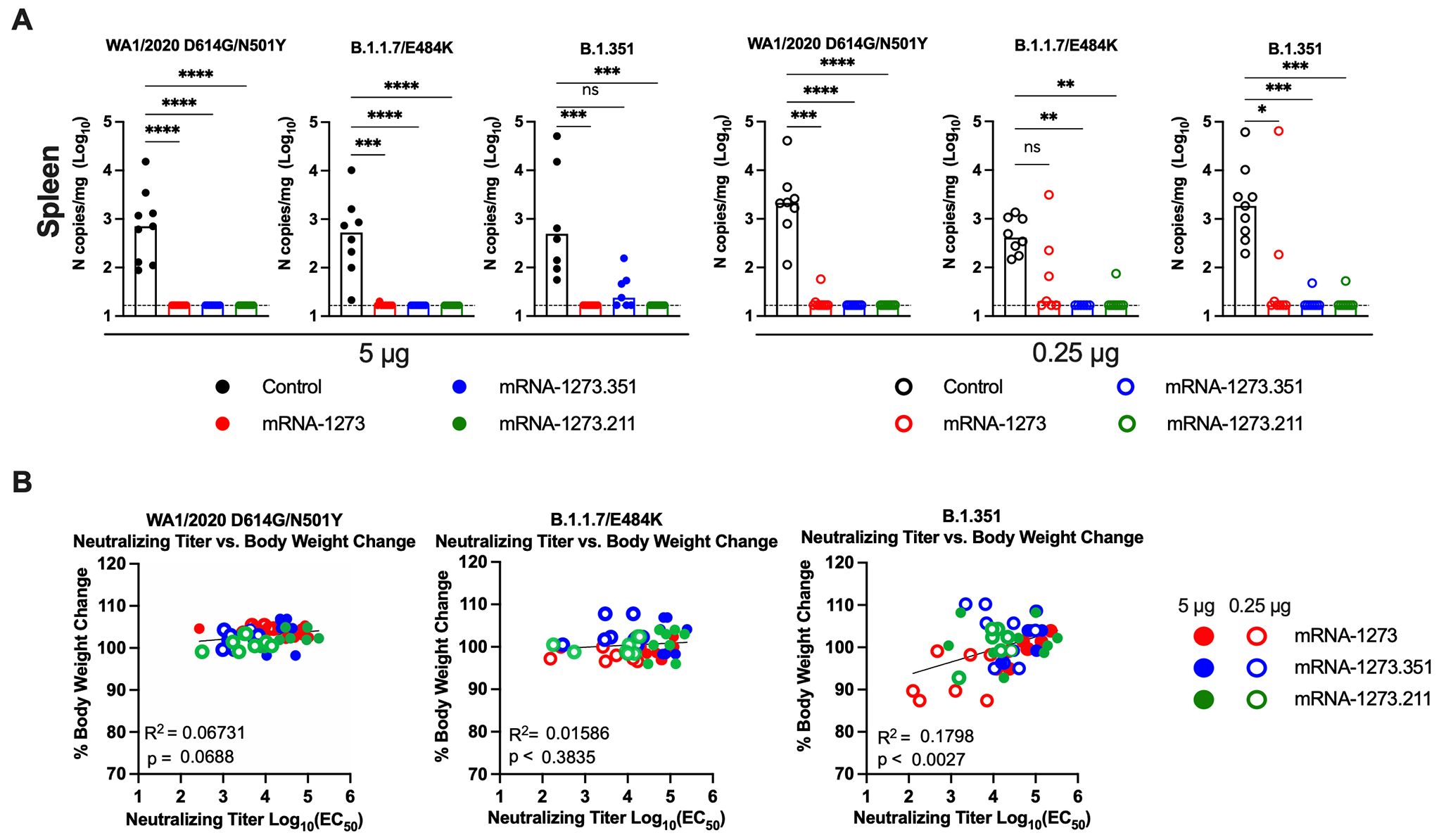

### Figure S3

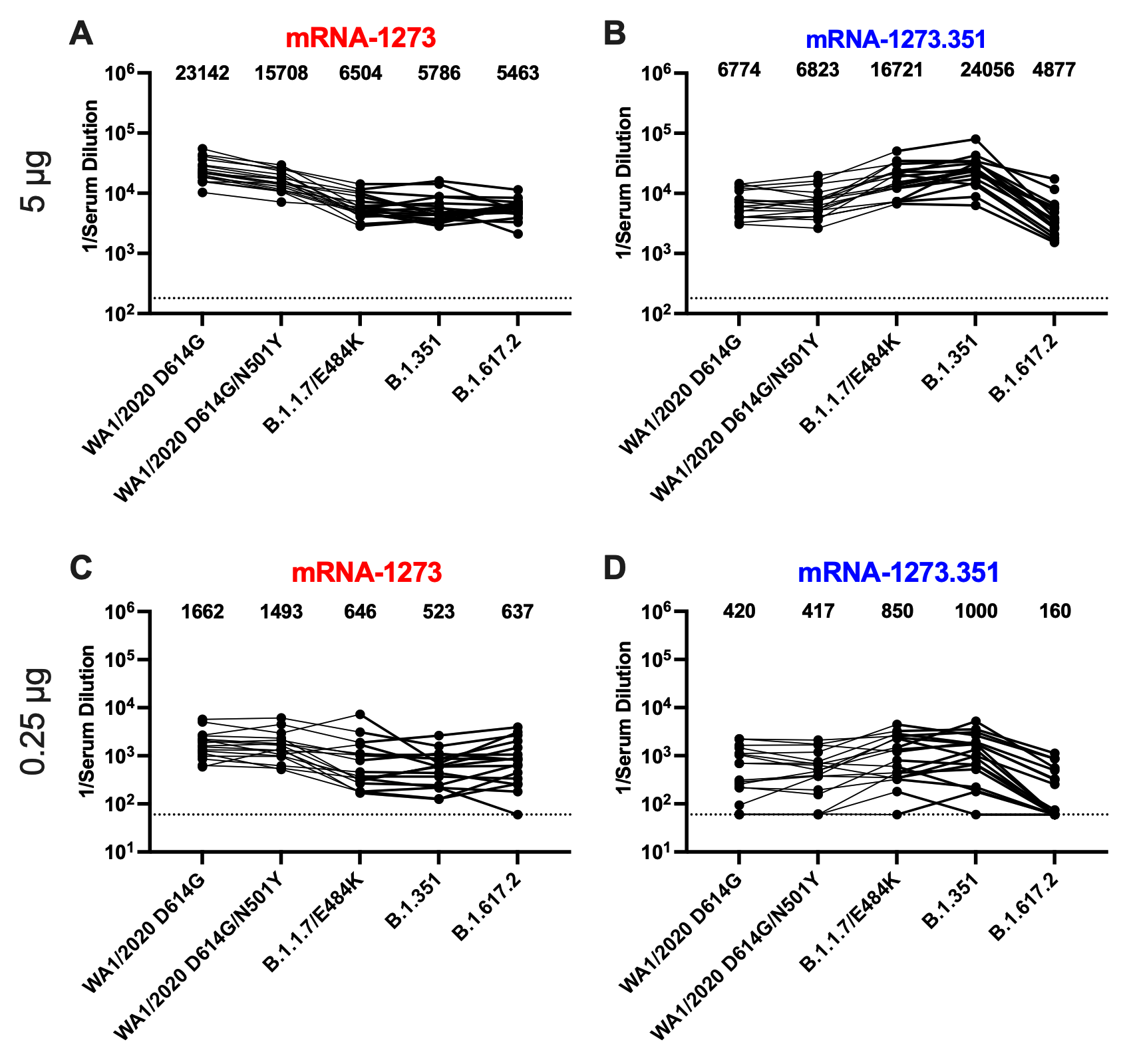

### Figure S4

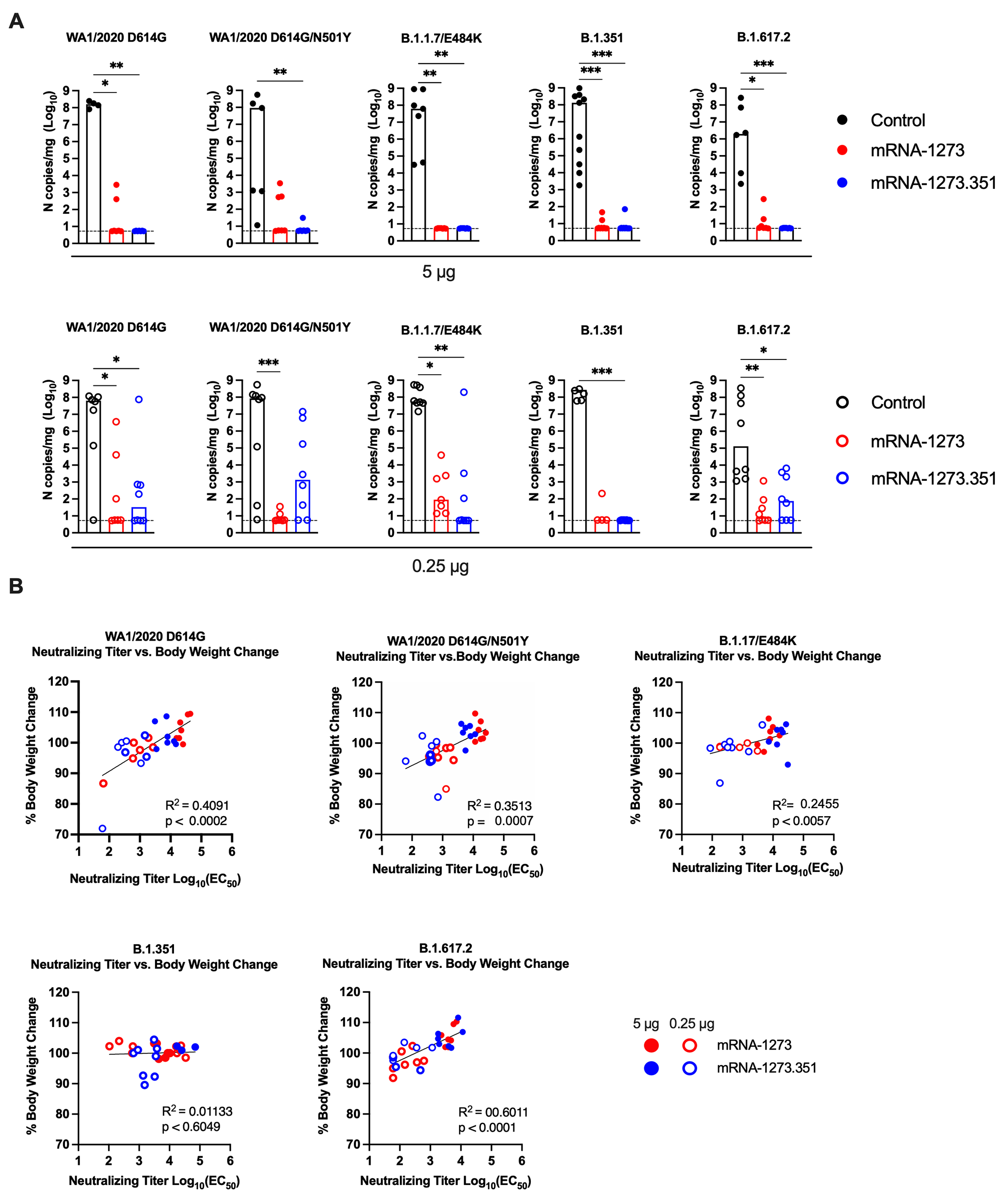

### Figure S5

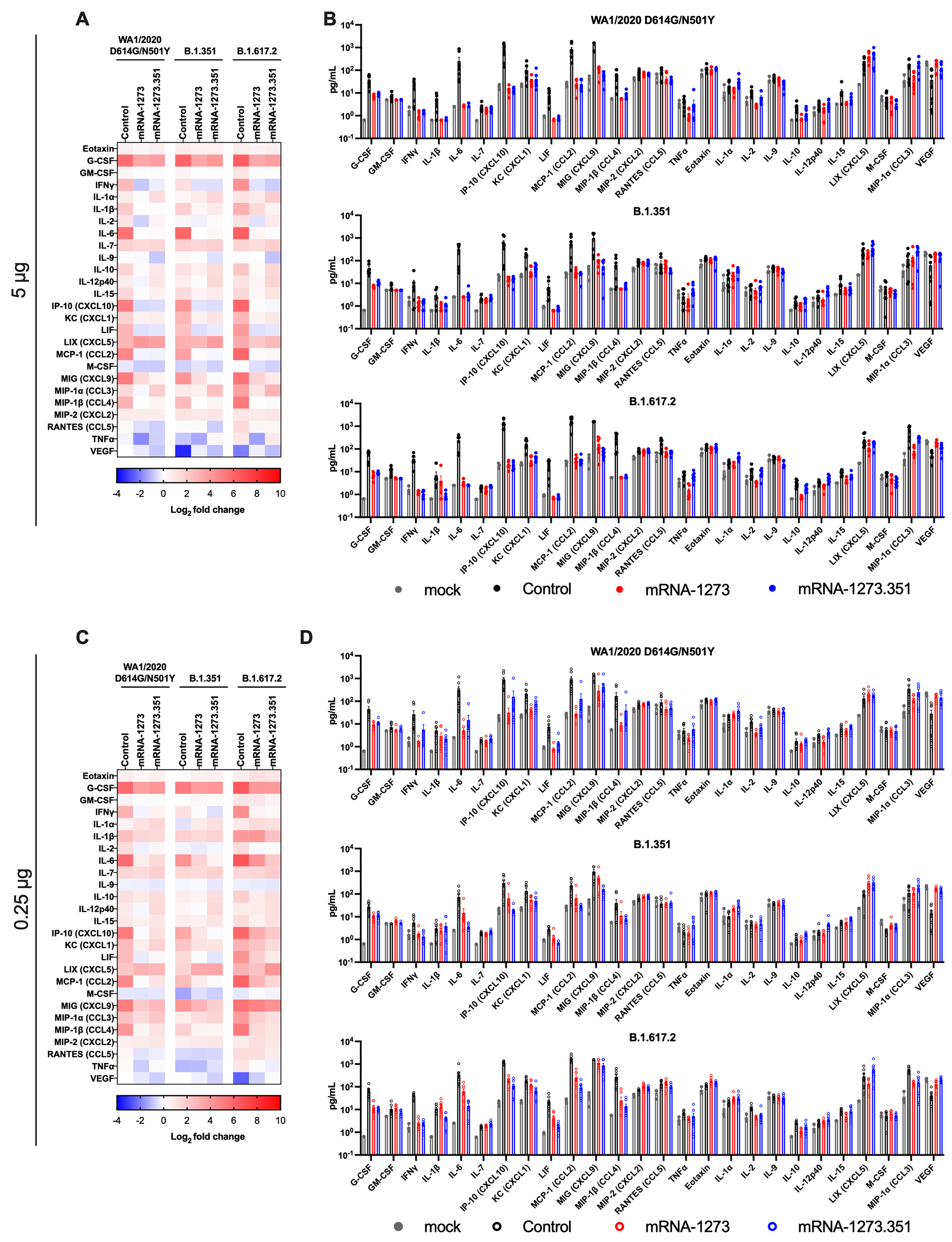
